# Systems modeling and uncertainty quantification of AMP-activated protein kinase signaling

**DOI:** 10.1101/2025.06.02.657503

**Authors:** Nathaniel Linden-Santangeli, Jin Zhang, Boris Kramer, Padmini Rangamani

## Abstract

AMP-activated protein kinase (AMPK) plays a key role in restoring cellular metabolic homeostasis after energy stress. Importantly, AMPK acts as a hub of metabolic signaling, integrating multiple inputs and acting on numerous downstream targets to activate catabolic processes and inhibit anabolic ones. Despite the importance of AMPK signaling, unlike other well-studied pathways, such as MAPK/ERK or NF-*κ*B, only a handful of mechanistic models of AMPK signaling have been developed. Epistemic uncertainty in the biochemical mechanism of AMPK activity, combined with the complexity of the AMPK pathway, makes model development particularly challenging. Here, we leveraged uncertainty quantification methods and recently developed AMPK biosensors to construct a new, data-informed model of AMPK signaling. Specifically, we applied Bayesian parameter estimation and model selection to ensure that model predictions and assumptions are well-constrained to measurements of AMPK activity using recently developed AMPK biosensors. As an application of the new model, we predicted AMPK activity in response to exercise-like stimuli. We found that AMPK acts as a time- and exercise-dependent integrator of its input. Our results highlight how uncertainty quantification can facilitate model development and address epistemic uncertainty in a complex signaling pathway, such as AMPK. This work shows the potential for future applications of uncertainty quantification in systems biology to drive new biological insights by incorporating state-of-the-art experimental data at all stages of model development.

## 1 Introduction

AMP-activated protein kinase (AMPK) is a master regulator of cellular metabolism that maintains energy homeostasis [1–6]. AMPK senses and responds to decreasing cellular energy availability [3, 4]. As a hub of metabolic signaling, AMPK integrates numerous inputs and acts on a large number of downstream targets [7–12]. Inputs to AMPK include cellular energy status [4, 8], intracellular calcium [13–15], and glucose levels [2]. AMPK is a heterotrimeric protein comprising an α-catalytic subunit and two regulatory subunits, the β- and γ-subunits [16, 17]. Phosphorylation of a key threonine residue, Thr172, on the γ-subunit leads to a 100-fold increase in the AMPK kinase activity and is catalyzed by many upstream kinases, such as LKB1 and CaMKK2 [5, 16]. Phosphorylated AMPK (pAMPK) is additionally dephosphorylated by phosphatases that include PP2A. Upon activation, AMPK phosphorylates a wide range of targets to promote catabolic processes and restore falling ATP levels [1, 3, 4], including acetyl-CoA carboxylase (ACC) [18], ULK1 [19], and RAPTOR [20]. The centrality of AMPK makes it a critical drug target [21], with well-known drugs including metformin [22, 23] and salicylate (Aspirin) [24] that act as AMPK activators.

A wide range of experimental approaches that include traditional molecular biology, modern omics, and structural biology have been used extensively to study the AMPK signaling network at the systems level [5, 7, 8, 25]. These studies have led to significant findings, such as the specificity of AMPK targets [8, 26], the mechanism of adenine nucleotide regulation of AMPK activity [16], and the *in vivo* response of AMPK to physiological stimuli such as exercise [25] (Figure 1). However, unlike other well-studied pathways such as MAPK/ERK [27, 28] or NF-*κ*B [29], which have 10s–100s of published mathematical models, fewer than five robust mechanistic models of AMPK signaling have been published [30, 31]. Several previous intracellular signaling models of metabolic pathways have embedded AMPK in a larger signaling network [9–11, 32]; however, these models often leverage phenomenological representations of AMPK activation as a function of select inputs and do not yet capture the regulation of AMPK activity in mechanistic detail. Alternatively, mechanistic models, such as those developed by Connolly et al. [31] and Coccimiglio et al. [30], aim to capture the details of AMPK activation and its interactions with adenine nucleotides in the contexts of neurotoxicity [31] and exercise [30]. While these previous models of AMPK signaling make well-justified assumptions, they were not rigorously constrained to high-quality AMPK signaling data from experiments. Here, we leveraged AMPK activity data measured using a recently developed fluorescent AMPK biosensor, ExRai-AMPKAR [33], to inform model development. These fluorescent biosensors are AMPK substrates and report relative phosphorylation by AMPK [33–35].

**Figure 1:**
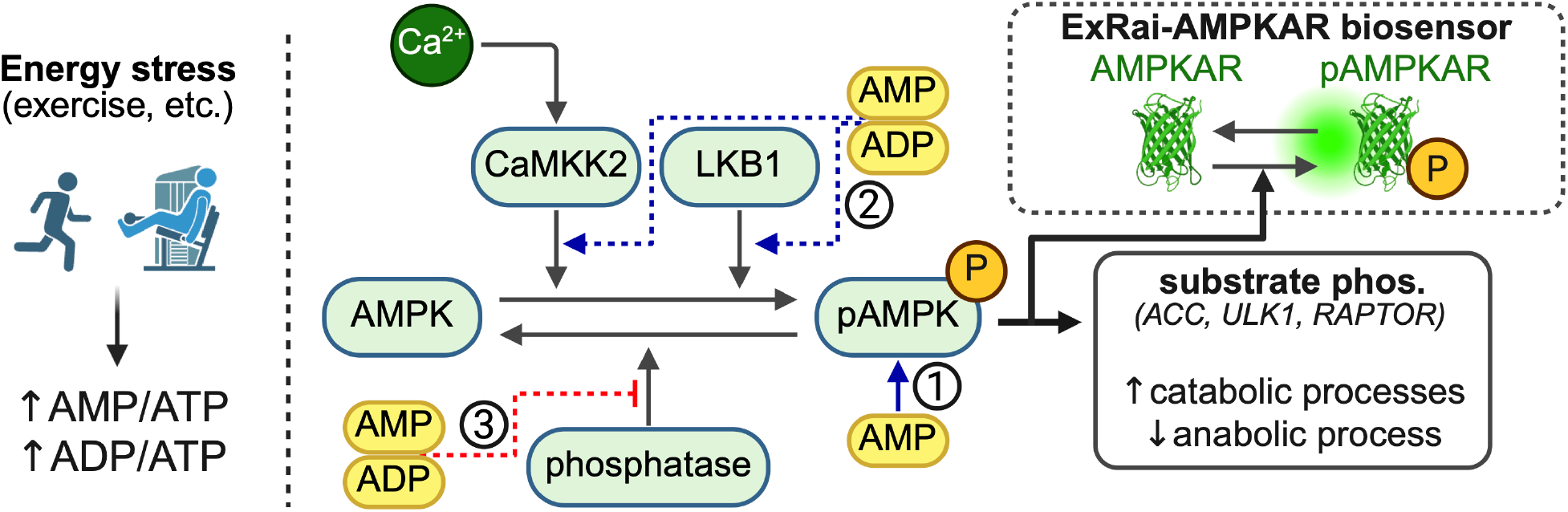
AMPK is activated by decreasing cellular energy availability that leads to falling intracellular ATP. Changes in intracellular adenine nucleotide concentrations, AMP/ATP and ADP/ATP, activate AMPK in three ways: (i) AMP acts as a direct allosteric activator of AMPK. (ii) AMP and ADP promote phosphorylation (activation) of AMPK by inducing conformational changes to the AMPK complex. (iii) AMP and ADP protect phosphorylated pAMPK from dephosphorylation. AMPK phosphorylates several downstream targets to promote catabolic and inhibit anabolic processes. AMPK activity can be measured using fluorescent kinase activity reporters such as ExRai-AMPKAR. These activity reporters are substrates of AMPK and report relative levels of phosphorylated by AMPK. Created in https://BioRender.com.

Model development for signaling networks typically leverages literature and domain expertise to identify relevant biochemical reactions and formulate a series of modeling assumptions [28, 36, 37]. These assumptions are then encoded into mathematical equations by prescribing kinetic terms, such as mass action or Michaelis-Menten kinetics, to represent reaction fluxes [38]. While such a workflow has led to numerous successful models, applying this approach to systems like AMPK, where new biology is still being discovered, can lead to a strong reliance on model assumptions. Recent research has introduced approaches for data-informed model development that leverage uncertainty quantification (UQ) [39] to ensure that models are well constrained to experimental measurements [40–42]. Here, we applied uncertainty quantification methods, including identifiability analysis [43], sensitivity analysis [44], Bayesian parameter estimation [40–42], and Bayesian model selection [28, 45, 46] to leverage AMPK activity data throughout the model development process.

In this work, we developed a set of related AMPK signaling models that vary in their mechanistic assumptions. Using Bayesian UQ, we estimated the associated kinetic parameters of these models using experimental data from Schmitt et al. [47], and selected the model that best captured the data. We then applied the new model to investigate AMPK signaling in the context of exercise. AMPK, which is thought to be activated during exercise, plays a multifaceted and complex role in regulating metabolism [48, 49] and modulating post-exercise adaptation [25, 50–52]. We therefore simulated different exercise regimes to generate physiologically relevant inputs to the AMPK signaling network. We found that AMPK is a robust integrator of high-frequency inputs, but shows exercise-dependent responses to longer timescale, low-frequency inputs. Our findings highlight how uncertainty quantification enables rigorous, data-informed predictive modeling for a system like AMPK, where our understanding of the biology is still evolving.

## 2 Methods and Materials

### 2.1 Workflow for model development with rigorous uncertainty quantification

Uncertainty quantification (UQ) aims to understand how errors in the data and models impact model predictions [39]. Recently, UQ methods for identifiability analysis [43, 53], sensitivity analysis [44],

parameter estimation [40, 54], and multimodel inference [28, 46, 55] have been applied in systems biology. Here, we leverage an end-to-end model development workflow to ensure that both model assumptions and predictions are constrained by available data (Figure 2). First, we developed a set of models that vary in the assumed mechanism and in the kinetic formulation used to represent the AMPK signaling system (Figure 2A). Next, we leveraged our previously developed approach for Bayesian parameter estimation that begins by reducing the dimensionality of the free parameter space using structural identifiability and global sensitivity analyses [40] (Figure 2B). In this work, we added the step of prior elicitation and refinement to ensure that parameters can only vary over regions of parameter space that lead to reasonable model predictions. Finally, we propagated uncertainty in the parameters forward to predictions by running ensembles of simulations, and used Bayesian model selection to choose the model that is most consistent with the data (Figure 2B). We summarize the details of each of these methods below. Supplemental Section S1.3 also highlights prior elicitation and refinement in more detail.

**Figure 2:**
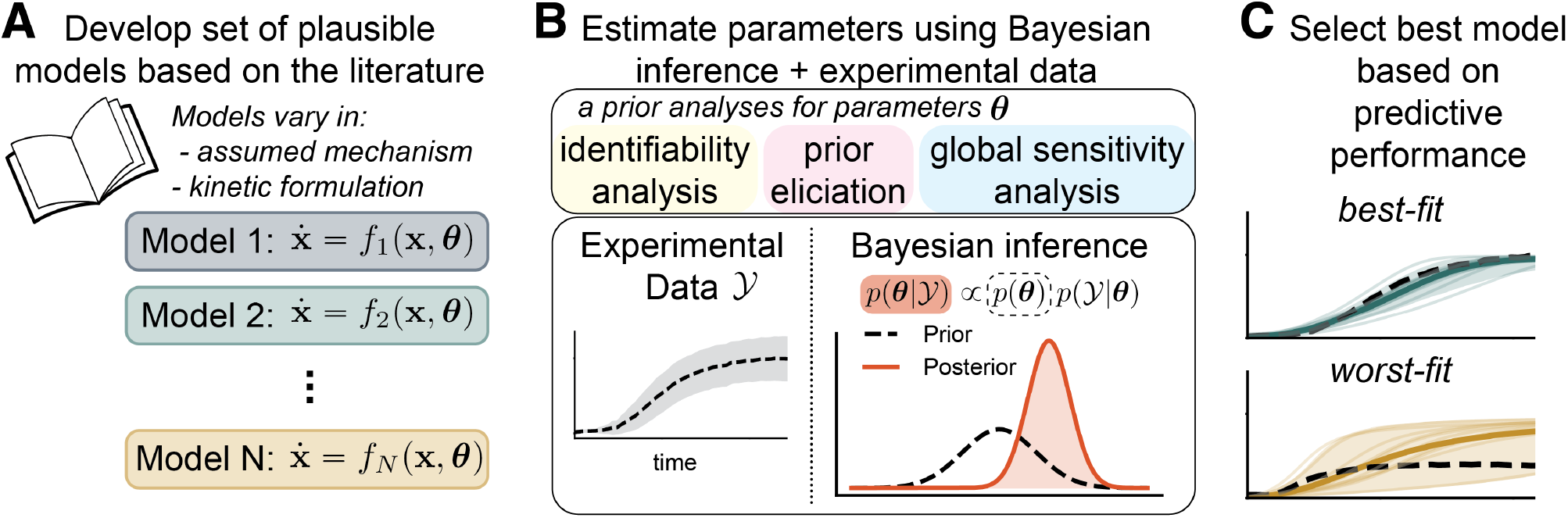
Uncertainty Quantification workflow used for data-informed model development. (**A**) First, we developed a set of AMPK signaling models based on available literature. The models vary in the assumed mechanism and the kinetic formulation. (**B**) Next, we estimated model parameters from experimental AMPK signaling data using Bayesian inference. We performed *a priori* identifiability and sensitivity analyses to reduce the dimension of the parameter space. Prior elicitation was used to ensure that priors lead to reasonable model predictions. (**C**) Finally, we used Bayesian model selection to select the best model based on predictive performance.

#### 2.1.1 Local structural identifiability analysis

Local structural identifiability implies that a parameter can be uniquely identified within a local region of parameter space space [43, 56]. In this work, we used the StructuralIdentifiability.jl package in the Julia programming language for local structural identifiability analysis [57]. Specifically, we used default settings for the `assess_local_identifiability()` function, and set the probability of correctness to *p* = 0.99 for all models. We fixed the parameters for the metabolism module and calcium-calmodulin-CaMKK2 reactions to nominal values. Supplemental Table 7 lists the identifiability of each model. We fix all nonidentifiable parameters to nominal values, which are potentially refined following the iterative process in Supplemental Section S1.3.

#### 2.1.2 Prior elicitation

We used direct prior elicitation [58] to find prior densities for free parameters such that 95% of the probability density falls between a specified range. Specifically, we solved an optimization problem for the prior hyperparameters that seeks to find the prior with the highest entropy that meets the 95% probability bounds using the Preliz Python library [59]. We specified the bounds to vary by several orders of magnitude around nominal values (Supplemental Table 1). We then ran prior predictive samples and performed prior predictive elicitation to refine the priors if the initial densities did not yield reasonable predictions [58, 59]. We detail this iterative refinement procedure in Supplemental Section S1.3. We used lognormal prior densities for all parameters.

#### 2.1.3 Global sensitivity analysis

Sobol global sensitivity analysis (GSA) partitions the total variance of an output quantity into the contributions of varying the uncertain model parameters over specified ranges [44]. Here we used the SALib Python library for GSA [60, 61]. Specifically, we allowed identifiable parameters to vary over ranges defined by the 90% highest-density interval of the prior density. Next, we generated a set of *N* random parameter samples from those ranges, where *N* is given by *N* = 256(*p* + 1) and *p* is the number of free parameters. We then simulated the models with these free parameters and calculated the maximum AMPKAR activation and time to half-maximum activation for the wild-type, LKB1 knockout, and CaMKK2 knockout conditions. Finally, we computed the Sobol total sensitivity index for each of the six quantities of interest (Supplemental Tables 8–13). We considered a parameter to be sensitive if it had a total sensitivity greater than 0.01 for any of the quantities. After this, the non-sensitive parameters were fixed to nominal values.

#### 2.1.4 Bayesian parameter estimation

In this section, we provide a brief overview of Bayesian parameter estimation. For more details in the context of systems biology, see [40, 54], and for general theory, see [39, 41]. We model the AMPK signaling network with a system of parametric ordinary differential equations (ODEs) that describe the rates of change of signaling species. Specifically, the models are defined as

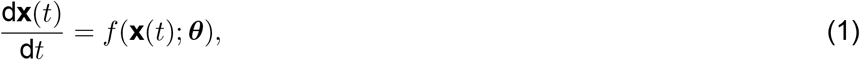

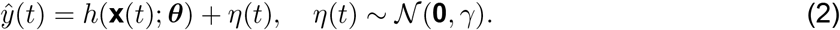

Here, the state variables, 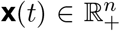, correspond to the concentration of biochemical species (ℝ_+_ = [0,∞) are the nonnegative real numbers). The observations, *ŷ*(*t*) ∈ [0, 1], are the fraction of activated AMPKAR biosensor. We assume that an independent and identically distributed Gaussian noise process, *η*(*t*) ∈ ℝ with variance *γ* ∈ ℝ. The function 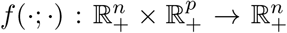 describes the system dynamics, and 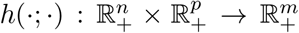 maps from states to observables. The model parameters 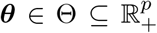 include reaction rates, equilibrium coefficients and other quantities that control model behavior.

Bayesian inference characterizes a probability density for the unknown model parameters conditioned on available data, p(***θ***|𝒴) [39, 41]. In this work, the training data, 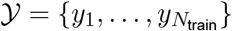 consists of *N*_train_ noisy experimental ExRai-AMPKAR observations. The probability density of the model parameters conditioned on data, called the posterior density, is learned using Bayes’ rule, which is defined as,

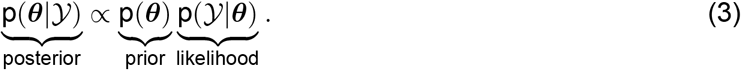

Here, p(***θ***) is the prior density which encodes assumptions about parameters before data is considered, and p(𝒴|***θ***) is the likelihood function that measures the probability that the model correctly predicts the training data. Because the models are nonlinear, we cannot evaluate the posterior density directly, so we instead must rely on methods such as *Markov chain Monte Carlo* (MCMC) [39–41] or *variational inference* (VI) [62] to characterize the posterior through the *S* samples drawn from it, {***θ***_1_, …, ***θ***_*S*_} ∼ p(***θ***|𝒴). In this work, we use the Pathfinder algorithm for VI [63], which approximates the posterior density with an easier-to-sample approximation by solving an optimization problem, because variational inference can be more computationally efficient than MCMC [62, 63]. We then draw samples from this approximating density. For a review of variational inference, see [62]. We implement all probabilistic models in the PyMC probabilistic programming language [64] and use default settings for the Pathfinder algorithm.

#### 2.1.5 Bayesian model selection

We use the expected log pointwise predictive density (ELPD) to assess predictive performance and direct model selection [41, 65]. Larger ELPD values indicate better predictive performance because they indicate a higher probability of correctly predicting out-of-sample data. We assume that the training data 𝒴 consist of statistically independent data points, such that 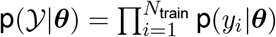. The ELPD is defined as

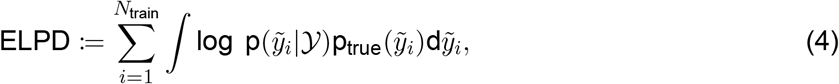

and quantifies the expected predictive performance of the model compared to the true data-generating distribution 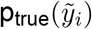, where 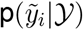 is the posterior predictive density of out-of-sample data point 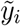. In general, the ELPD is computationally intractable because we do not know the true data-generating density, 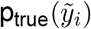, so instead we approximate the ELPD using the leave-one-out cross-validation estimator (LOO-CV) [65]. Specifically, we use the ArviZ Python library [66], which uses Pareto-smoothed importance sampling to compute the LOO-CV estimator [67]. We use default settings for all computations.

### 2.2 ODE simulation

We solved all differential equations numerically using the Kvaerno 4/5-order implicit Runge Kutta method, which is implemented in the Diffrax Python library [68, 69]. Additionally, we used a PID controller-based adaptive time-stepping algorithm with tolerances `atol=1e-6` and `rtol=1e-6` [70, 71]. We chose the Diffrax library because it enables us to compute gradients of the ODE solution with respect to the model parameters using autodifferentiation, which is necessary for employing Pathfinder VI for parameter estimation.

### 2.3 Experimental data pre-processing

The ExRai-AMPKAR AMPK kinase activity reporter, developed in [47], is a ratiometric biosensor in which the ratio of emission intensities at two excitation wavelengths increases as the sensor is phosphorylated by active AMPK. Therefore, the emission intensity ratio is considered to vary with the degree of AMPK kinase activity. We aimed to calibrate the ratiometric readout to the fraction of phosphorylated sensor, which can be computed from model outputs. Similar normalizations are reviewed in [72, 73]. To normalize the data, we assumed that (i) the concentration of phosphorylated ExRai-AMPKAR is negligible before 2-DG stimulation, (ii) nearly all expressed ExRai-AMPKAR is phosphorylated when the ratiometric signal is at its maximum, and (iii) the fraction of phosphorylated sensor varies proportionally to the ratiometric signal. Based on these assumptions, we renormalized the average normalized ratiometric AMPKAR data, defined as *R*(*t*), using the normalization

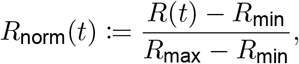

where *R*_min_ and *R*_max_ are the minima and maxima of the signal in time. We additionally normalized the variance of the signaling by the same normalization factor. We used the minimum and maximum signals from the wild-type condition to normalize the kinase knockout conditions. To predict the renormalized signals from our models, we assume that

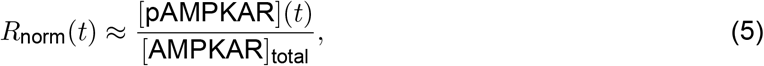

where [pAMPKAR](*t*) is the time-varying concentration of the phosphorylated biosensor, and [AMPKAR]_total_ is the total concentration of both active and inactive ExRai-AMPKAR. We drop “ExRai-” for brevity.

### 2.4 Statistical comparison

We use the pairwise Student’s t-tests with the Bonferroni correction to adjust for multiple comparisons to test for statistical significance of the difference between population means. We assess significance at a level of *α* = 0.05. To effectively summarize comparisons between multiple groups, we employ a compact letter display, where groups with the same letter do not exhibit statistically significant differences in their means.

## 3 Results

### 3.1 Model development

To capture AMPK activation in response to energy stresses, we developed a set of six mechanistic models of the AMPK signaling pathway (Figure 1) using the UQ workflow outlined in Figure 2. We included a minimal representation of metabolic processes, including glycolysis, oxidative phosphorylation, and ATP hydrolysis, and captured key mechanisms of AMPK activation and regulation, including AMPK phosphorylation and interactions with adenine nucleotides [1, 5]. Although we used previous models, including [9, 30], during model development, our new models differ in several ways. First, we included the specific mechanisms by which adenine nucleotides regulate AMPK activity in our models. Second, we explicitly included the upstream kinases, LKB1 and CaMKK2, to capture AMPK activation due to energy stress and calcium signaling. These kinases are known to activate AMPK via phosphorylation [13–15, 74]. Third, we varied the associated mechanisms between the six models to investigate how adenine nucleotide binding to AMPK affects AMPK phosphorylation and activity. Importantly, we investigated models that employ different kinetic formulations for the reaction rates before constructing models of AMPK signaling. The remainder of this section delineates key model assumptions and describes our models.

#### 3.1.1 Kinetic formulations impact parameter identifiability and predictive accuracy

Models of intracellular signaling typically prescribe kinetic equations, including mass action, Michaelis-Menten, and Hill-type, to describe the dynamics of biochemical reactions [36, 38]. If the reaction mechanism is well known, then the best practice is to choose the corresponding kinetic formulation, for example, prescribing Hill-type kinetics when a reaction is cooperative [38]. However, when little is known about a reaction mechanism, kinetic formulations are often prescribed in an *ad hoc* or phenomenological fashion to ensure that a model retains the flexibility to capture available experimental observations [36]. However, if these modeling choices are made while considering impacts on parameter and predictive uncertainties, then subsequent parameter estimation will be more successful [40].

To understand how different kinetic formulations impact parameter estimation and model predictions, we first modeled a single enzyme-catalyzed reaction using mass action, Michaelis-Menten, and Hill-type kinetics. The reaction

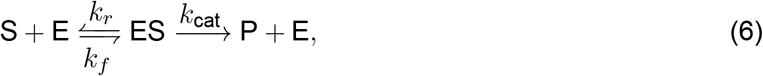

describes the formation of product P from substrate S, catalyzed by conserved enzyme E. The reaction rates *k*_*f*_, *k*_*r*_, and *k*_cat_ describe the rate of substrate-enzyme binding and unbinding, and the rate of formation of the product, respectively. In Figure 3 we show the three models; we note that the number of equations and free parameters varied between the models. Interestingly, the local identifiability of the model parameters was different for these different reaction rate formulations (Figure 3). We found that all model parameters for the three models are locally identifiable when the product concentration is observed directly, *y*_obs_ = [*P*]. However, the identifiability differed between the model formulations when the ratio of the concentration of product to the total concentration of substrate, *y*_obs_ = [*P*]/[*S*]_tot_, was observed. Notably, two of the three parameters of the mass action model were locally identifiable, while none of the Michaelis-Menten or the Hill-type models were identifiable.

**Figure 3:**
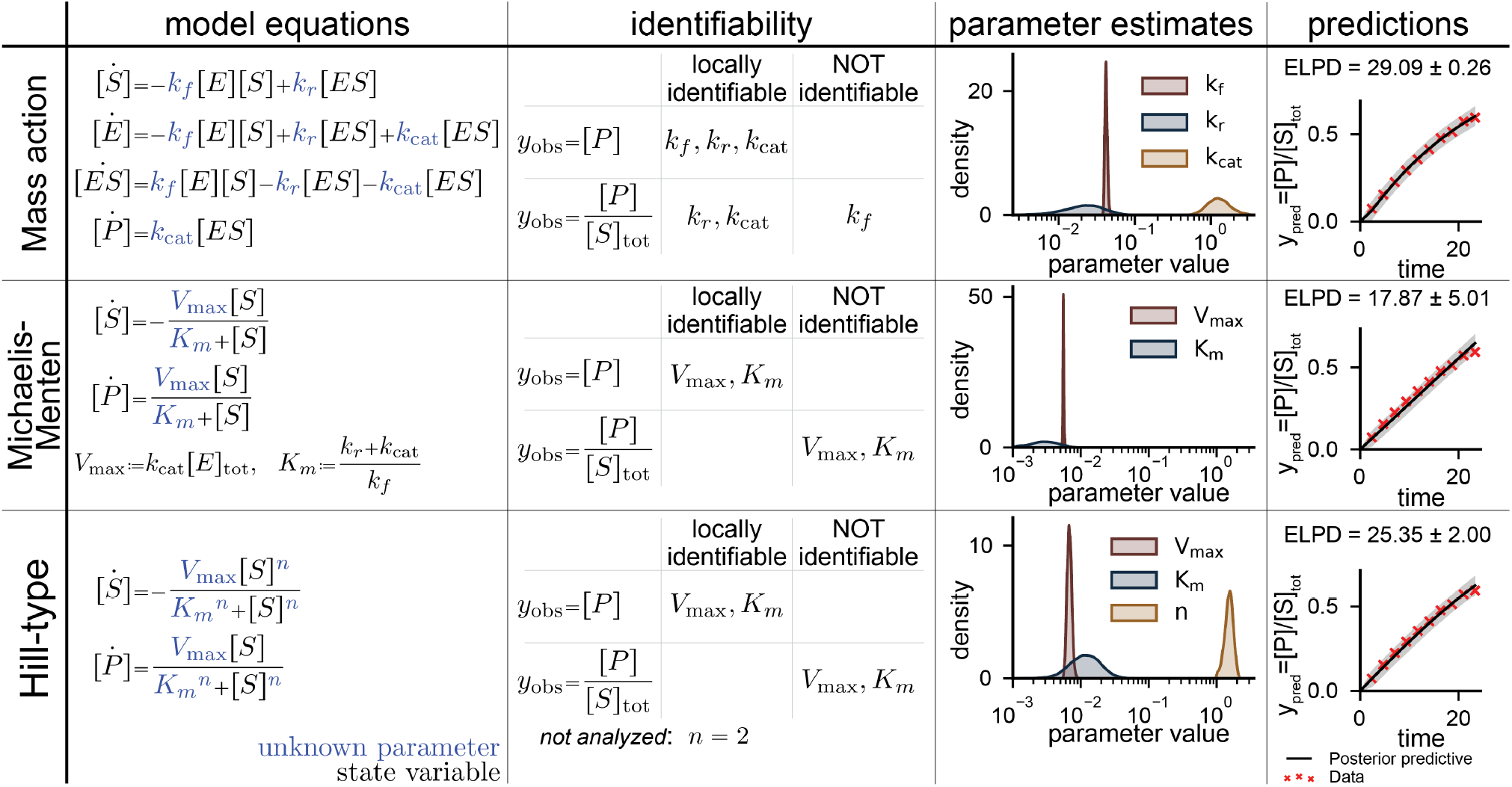
Identifiability varies between mass action, Michaelis-Menten, and Hill-type models of a single enzyme-catalyzed reaction. *Model equations*: mass action, Michaelis-Menten, and Hill-type kinetics can be used to formulate differential equation models of a single enzyme-catalyzed reaction. Uncertain free parameters are shown in blue. *Identifiability:* Different kinetic formulations yield different structural local identifiability from direct observations of the concentration of the product (*y*_obs_ = [*P*]) and more experimentally feasible observations of the ratio of product to total substrate 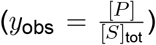. *Parameter estimates:* All free model parameters are estimated from data of the ratio of product to total substrate shown in red. Black traces show the posterior mean and gray shading shows the 95% posterior credible interval computed using 8,000 posterior samples.

Next, to assess how structural nonidentifiabilities impacted parameter estimation, we fit the three models to experimental data of the phosphorylation reaction catalyzed by pyruvate kinase from [75]. In the reaction, phosphoenolpyruvate is converted to pyruvate by the pyruvate kinase enzyme, which, under the experimental conditions, can be considered a single-substrate, single-enzyme reaction. We estimated the distribution of all model parameters using Bayesian inference to gain a better understanding of how nonidentifiabilities affect the estimates. Specifically, we normalized the data to a total substrate concentration of 0.195 mM used in the experiments [75] to mimic the data that observes the product as a fraction of the total substrate. In each of the models, we were able to estimate certain parameters with high certainty (*k*_*f*_ and *V*_max_) and others with low certainty (*k*_*r*_ and *K*_*m*_; Figure 3). Next, we simulated the models using the posterior parameter samples to propagate the parametric uncertainty forward to predictions (Figure 3). The mass action model had the highest probability of correctly predicting future data measured by the expected log pointwise predictive density (ELPD; see Eq (4) for the definition; ELPD = 29.09). The Hill-type model was next best (ELPD = 25.35), and the Michaelis-Menten model was the worst of the three (ELPD = 17.87).

Since systems biology models typically involve cascades of coupled biochemical reactions, we constructed nine models of the two-reaction system in which a substrate is phosphorylated by a kinase to form a product, which is then dephosphorylated (Supplemental Section S1.1). Interestingly, we found that structural identifiability of the model parameters is not impacted by the coupling of two reactions (Supplemental Table 3). For example, the identifiability of the model parameters for the model that uses mass action kinetics for both the phosphorylation and dephosphorylation reactions (Supplemental Table 3) was the same as the identifiability for the single reaction mass action model (Figure 3).

Overall, for the single-reaction system, the three model formulations were all limited in their ability to capture the pyruvate kinase reaction without introducing structural nonidentifiabilities or increasing the number of equations and parameters in the model. The mass action formulation appeared to be a safe choice because it had the most favorable identifiability, and the additional reaction (and equation) did not introduce a significant computational burden. Contrary to common belief, the Michaelis-Menten formulation did not seem to be the best choice for this example, because it had the lowest predictive accuracy (lowest ELPD). The Hill-type model was almost as good as the mass action model in terms of predictive performance; however, we found that the inclusion of the Hill coefficient, *n*, made parameter estimation more difficult [40]. Therefore, we concluded that for these simple reactions, the mass action models had more favorable identifiability and yielded higher-quality predictions. However, despite the nonidentifiabilities, we could still estimate the parameters of the Michaelis-Menten and Hill-type models.

#### 3.1.2 Proposed AMPK signaling models describe a range of regulation and signaling mechanisms

Since we did not find a *best* kinetic formulation for describing reaction rates in terms of identifiability in the previous section, we built six related models that use either mass action or Michaelis-Menten kinetics. Additionally, the models differed in their AMPK regulatory mechanisms. Table 1 highlights the differences between the models. Each model consisted of three core modules: (i) cellular energetics and metabolism, (ii) AMPK regulation, and (iii) AMPK kinase activity and biosensor. In this section, we outline our assumptions and summarize the three core modules. We provide the model equations and initial conditions in the Systems Biology Markup Language (SBML) format, as well as Python functions, in the available code. We list nominal parameters in Supplemental Tables 5–6.

**Table 1:** Set of AMPK models vary in assumed reaction mechanisms.

| Model Number | Kinetics | Allosteric activation by AMP | Promotion of phosphorylation | Protection from dephosphorylation | Antagonism by ATP |
| --- | --- | --- | --- | --- | --- |
| 1 | mass action (MA) | AMP-pAMPK is the only active kinase | only AMP/ADP-AMPK are phosphorylated by LKB1 | AMP-pAMPK and ADP-pAMPK cannot be dephosphorylated | ATP-AMPK not phosphorylated by LKB1 & ATP-pAMPK not active |
| 2 | Michaelis-Menten (MM) | same as 1 | same as 1 | same as 1 | same as 1 |
| 3 | MA | AMP is a nonessential act. (activity enhanced by $\beta_{\text{AMP}} \geq 1$) | AMP/ADP reduce LKB1-AMPK $K_d$ by $\alpha_{\text{LKB1}} < 1$ | AMP/ADP increase PP-pAMPK $K_d$ by $\alpha_{\text{PP}} > 1$ | same as 1 |
| 4 | MM | same as 3 | same as 3 | same as 3 | same as 1 |
| 5 | MM | AMP is a nonessential act. (activity enhanced by $\beta_{\text{AMP}} \geq 1$) | AMP/ADP increase LKB1 phosphorylation rate by $\beta_{\text{LKB1}} > 1$ | AMP/ADP decrease PP phosphorylation rate by $\beta_{\text{PP}} < 1$ | n/a |
| 6 | MM | same as 5 | same as 5 | same as 5 | same as 5 |

##### Module 1: Cellular metabolism and energy stress

AMPK senses changes in the cellular energy state through changes in adenine nucleotide levels [1, 5]. Cellular metabolism is a highly regulated process that tightly controls the production of ATP and the conversion between ADP and AMP [3]. To ensure that our models maintain physiologically consistent adenine nucleotide levels, we adapted the simplified metabolism models from Coccimiglio et al. [30] and Lueng et al. [9] to model cellular ATP production, consumption, and conversion between AMP, ADP, and ATP. We modeled ATP generation by both oxidative phosphorylation and glycolysis, and ATP consumption via a global hydrolysis reaction. Additionally, we included the adenylate cyclase and the creatine kinase reactions to ensure that the appropriate stoichiometry is maintained between adenine nucleotides. Throughout this work, we assumed that the initial concentrations of AMP, ADP, and ATP are 7.03 mM, 1.11 mM, and 7.9 × 10^−2^ mM, respectively, which we found to be a steady state of the metabolism model (Supplemental Figure S1A). We modeled energy stresses due to the 2-deoxyglucose (2-DG) stimulus applied in [47] by decreasing the rate of glycolytic ATP production 100-fold (Supplemental Figure S1B–D). Additionally, we modeled energy stress due to increased ATP consumption, such as during exercise, by increasing the ATP hydrolysis rate *k*_hydro_ or by prescribing an additional negative ATP flux. We used the same metabolism module across all the models that we developed. Supplemental Section S1.2.1 describes the metabolism model in more detail, and Supplemental Tables 4–5 provide the flux equations and reaction parameters, respectively.

##### Module 2: Adenine nucleotide binding and AMPK phosphorylation

AMPK activity is regulated by changes in adenine nucleotide concentrations through a tripartite mechanism that modulates net AMPK phosphorylation and directly activates AMPK (Figure 1) [1, 3, 5, 16, 17]. Net AMPK phosphorylation is regulated by several effects of AMP and ADP binding that induce conformational changes to the AMPK complex, including promoting phosphorylation at Thr172 and protecting from dephosphorylation (Figure 1). Adenine nucleotides can bind to four cystathionine-βsynthase (CBS) repeats on the γ-subunit of AMPK [5, 16]. However, recent evidence suggests that only binding in the third CBS repeat regulates AMPK activity [5]. Thus, we only accounted for single adenine nucleotide-AMPK binding interactions and assumed that adenine nucleotides can bind and unbind from AMPK.

We modeled each of the three parts of the tripartite AMPK regulatory mechanism (Figure 1). First, AMP allosterically activates AMPK, in which AMP-AMPK binding has been found to result in a 10-fold activation in AMPK kinase activity [16, 18, 76–78]. Second, AMP and possibly ADP binding have been found to promote AMPK phosphorylation by upstream kinases by inducing conformational changes that expose Thr172 to these kinases [16, 79, 80]. Third, AMP and ADP binding protect Thr172 from dephosphorylation [16, 76, 78]. Notably, ATP is thought to antagonize the effects by competing for the third CBS binding site and induce inhibitory effects on AMPK [5, 17, 21]. Thus, AMPK is sensitive to the AMP/ATP and ADP/ATP ratios.

We constructed three sets of models that make distinct assumptions about the mechanism of adenine nucleotide binding interactions, all of which are potentially valid according to recent literature. In our models, LKB1 activity remains constant, and only interactions between adenine nucleotides and AMPK regulate net LKB1-mediated phosphorylation. We assumed that AMP and ADP are essential activators of LKB1 in Models 1 and 2, where AMP or ADP binding is necessary for LKB1 to phosphorylate AMPK [38]. Alternatively, in Models 3–6, we assumed that AMP and ADP are nonessential activators of LKB1, where AMP and ADP binding either decrease the *K*_*d*_ of LKB1 binding (Models 3 and 4) or increase the rate at which LKB1 phosphorylates the AMPK complex (Models 5 and 5) [38]. In addition to LKB1, we included CaMKK2, which is known to phosphorylate AMPK in a Ca^2+^-dependent manner [14, 24]. We assumed that CaMKK2 activity depends on the intracellular Ca^2+^ concentration through calcium-calmodulin binding with the kinetics from [81, 82]. In Models 1–4, we assumed that adenine nucleotide binding does not affect CaMKK2-mediated phosphorylation, such that all AMPK complexes are phosphorylated with the same kinetics, because there has been debate as to whether CaMKK2 activity is impacted by adenine nucleotide binding [5, 8, 16, 21, 76]. However, to capture the possibility that AMP and ADP promote CaMKK2-, in addition to LKB1-, mediated AMPK phosphorylation, in Models 5 and 6, we modeled these reactions with a nonessential activation mechanism. Lastly, we modeled the net dephosphorylation of phosphorylated pAMPK with a single dephosphorylation reaction, assuming that AMP and ADP, but not ATP, protect from dephosphorylation. In Models 1 and 2, we assumed that AMP and ADP binding completely protect pAMPK from dephosphorylation while free pAMPK and ATP-bound pAMPK are susceptible to dephosphorylation. In Models 3–6, we assumed that AMP and ADP binding reduce the effectiveness of pAMPK dephosphorylation via nonessential inhibition [38]. We assumed the total enzyme concentrations and initial conditions are the same across all six models.

##### Module 3: AMPK kinase activity and fluorescent biosensor

Phosphorylated AMPK is an active kinase with many downstream targets [7]. As its name suggests, AMPK is activated nearly 10-fold by AMP binding [16, 18, 76–78]. Here, we modeled AMP as either an essential activator of AMPK (Models 1 and 2) or a nonessential activator of AMPK (Models 3–6). To enable direct calibration of our models to fluorescent microscopy data of AMPK activity from [47], we explicitly modeled the AMPK activity biosensor, ExRai-AMPKAR, as an AMPK substrate [72, 73]. We refer to the ExRai-AMPKAR biosensor as AMPKAR for brevity. In all the models, we assumed that active AMPK phosphorylates AMPKAR [34, 35, 47]. Additionally, we modeled net phosphorylated pAMPKAR dephosphorylation with a single dephosphorylation reaction. We assumed that the fraction of pAMPKAR to total AMPKAR is directly equivalent to the fraction of the above baseline fluorescent sensor response (Eq (5)).

### 3.2 Application of the UQ framework suggests that AMP and ADP act as nonessential activators of AMPK

Having developed a set of AMPK signaling models that reflect possible AMPK regulatory mechanisms, we applied the UQ-based model development workflow shown in Figure 2. First, we estimated model parameters to constrain the model predictions to experimental data. Although there are some direct experimental observations of the activities of LKB1, CaMKK2, AMPK, and the related phosphatases, and with AMPK adenine nucleotide binding affinities, these data were not collected in the same cellular context as the available experimental data [16, 74, 77, 83]. Therefore, we aimed to estimate unknown model parameters using physiologically relevant data of energy-stress induced AMPK activity [47]. The number of free parameters ranged from 16 to 25 Model 1 and Model 5, respectively. Here, we used Bayesian parameter estimation because it allows us to leverage previous knowledge of parameters to inform the parameter estimates and to quantify uncertainty in estimates [39–41]. Bayesian estimation represents unknown parameters as random variables and aims to infer their probability densities conditioned on experimental data and prior knowledge (see Section 2.1.4).

We have previously found that restricting the set of estimated parameters to identifiable and influential parameters is necessary for successful parameter estimation [40]. So, we performed structural identifiability and global sensitivity analyses before estimating any parameter (Supplemental Table 7). For the structural identifiability analysis, we fixed the metabolism and calcium-calmodulin model parameters to the nominal values listed in Supplemental Table 1, because we had well-established models for those components [9, 30, 81, 82]. Interestingly, contrary to the results for the single-enzyme reactions, we found that all of the mass action model parameters (Models 1, 3, and 5) were structurally locally identifiable. Most of the Michael-Menten model parameters (Models 2, 5, and 6) were identifiable, except for those that control AMPK phosphorylation of the sensor and sensor dephosphorylation. For the subsequent analyses and parameter estimation, we fixed nonidentifiable parameters to the nominal values listed in Supplemental Table 1.

Next, we performed global sensitivity analysis using Sobol sensitivity analysis to determine which uncertain parameters have the greatest influence on model predictions in response to a severe energy-stress-like stimulus [44]. We specified the plausible ranges for parameters spanned by several orders of magnitude around the nominal parameters (Supplemental Table 1). However, as these ranges were loosely based on experimental evidence, we utilized prior predictive simulations to determine if the chosen parameter ranges and nominal values for nonidentifiable parameters were reasonable [41, 58] (see Section 2.1.2). We found that initial parameter ranges and the nominal values for nonidentifiable parameters did not necessarily lead to reasonable predictions (Supplemental Figure S3). For example, for Model 1, the distribution of initial values was, as expected, mostly centered around zero (Figure 4A); however, although the distribution of final values was well spread between zero and one, most of the predictions reached the maximum value too quickly (Figure 4B and 4C). We hypothesized that a combination of unfeasible parameter ranges for the free parameters and incorrect nominal values for the fixed nonidentifiable parameters was responsible for the poor predictions. Therefore, we took an iterative approach to refine the parameter ranges and nominal values such that the prior predictive density of predictions was reasonable (Supplemental Section S1.3 and Supplemental Figures S2 and S3). After refining the parameter ranges, we analyzed the total sensitivity of the maximum predicted AMPK sensor response and the time to half-maximum for three scenarios: (i) wild-type kinase activity, (ii) LKB1 knockout in which only CaMKK2 is active, and (iii) CaMKK2 knockout (Supplemental Tables 8–13). We found that most free parameters influenced at least one of the quantities since they have a total Sobol sensitivity greater than 0.01. We fixed any parameters not meeting this threshold to their corresponding nominal values and left the remaining parameters free for parameter estimation.

**Figure 4:**
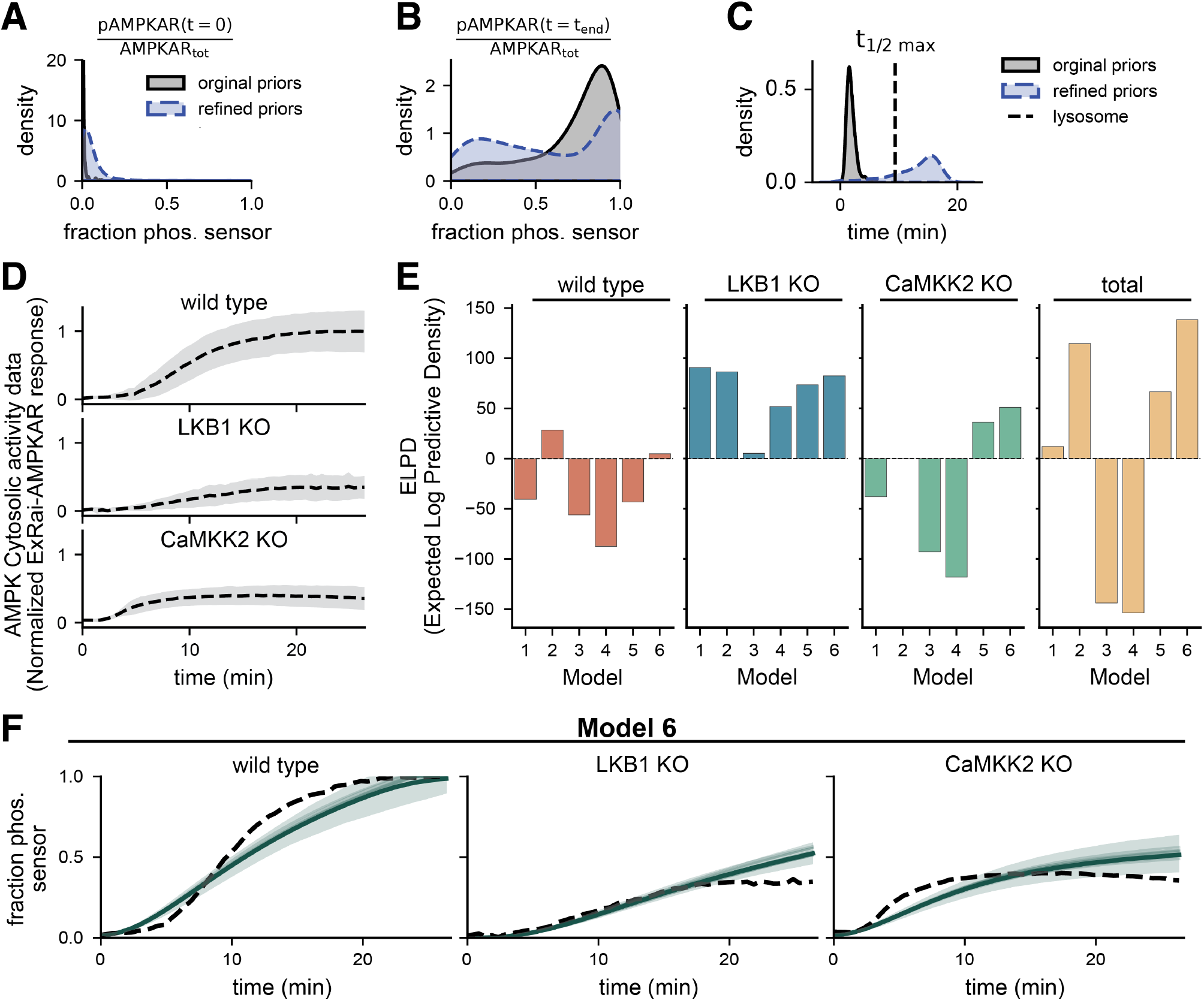
UQ workflow selects models that treat AMP and ADP as nonessential activators of AMPK. (**A**-**C**) Estimated probability densities of key quantities-of-interest of the prior predictive density of the mass action model before (black) and after (blue) refining priors with predictive prior elicitation. Priors were refined to ensure the densities placed probability mass on desired values. Supplemental Figure S3 shows samples from the prior predictive distribution that were used to generate densities of key quantities. (**D**) Experimental data of AMPK activity in response to 2-deoxyglucose (2-DG) induced energy stress. Data shows the normalized ExRai-AMPKAR signal, which captures AMPK activity by varying proportionally to the fraction of biosensor phosphorylated by upstream AMPK. Experimental conditions include wild type, LKB1 knockout mutants (LKB1 KO), and CaMKK2 knockout mutants (CaMKK2 KO). Data originally from [47]. (**E**) Predictive performance is quantified by the expected log pointwise predictive density (ELPD) across conditions in the data. Higher ELPD values indicate better predictive performance and lower uncertainty. Models were calibrated to data from all three experimental conditions. (**F**) Posterior predictions generated using 1000 posterior samples from Model 6, which had the highest total ELPD. The dashed black line shows the data for each condition. The solid green line shows the posterior mean, and the shaded band shows the 95% posterior credible interval.

After determining which parameters are identifiable and influential, we used Bayesian inference to estimate probability densities for the uncertain parameters conditioned on fluorescence microscopy AMPK activity data from Schmitt et al. [47]. We used the recordings of cytosolic AMPK activity (Figure 4A) for parameter estimation, and we simulated the 2-deoxyglucose (2-DG) stimulus from experiments by decreasing the rate of ATP production via glycolysis 100-fold from 0.5 1/s to 0.005 1/s, because 2-DG blocks glycolysis. Initially, we aimed to estimate model parameters from the wild-type data alone; however, we found that this data did not inform the individual LKB1 and CaMKK2 activities, so the calibrated models could not accurately predict the kinase knockout conditions (Supplemental Figure S4). Therefore, we used the data from all three conditions for parameter estimation.

After successfully estimating posterior densities for the parameters (Supplemental Figures S5 and S6), we used Bayesian model selection to determine which reaction mechanism and kinetic formulation best capture the data [28, 45, 65]. Specifically, we computed the expected log pointwise predictive density (ELPD; defined in Eq. (4)) of each model, which quantifies the probability of correctly predicting unseen data [45, 65]. Higher ELPD values indicate better predictive performance. We found that only Model 6 could predict all three conditions well, so the total ELPD for that model was the greatest (total ELPD=138.25; Figure 4E). The next best model, Model 2 (total ELPD=66.48), could not capture the CaMKK2 knockout condition, which suggests that the essential activation mechanism of LKB1-mediated phosphorylation is incorrect in that model. Furthermore, Model 5, which used mass action kinetics and had similar nonessential mechanisms to Model 6, showed poorer predictions than Model 6. Figure 4F shows the posterior predictions using Model 6. These results suggest that AMP and ADP act as nonessential activators of AMPK, increasing net AMPK phosphorylation and AMPK kinase activity. Furthermore, in general, the models that prescribed the Michaelis-Menten kinetics performed better than those that used mass action. Based on these results, we used Model 6 for subsequent simulations.

### 3.3 AMPK activity is sensitive to the timing of input stimuli

Next, we sought to understand how stimulus strength and timing affect AMPK activity. We previously predicted the fraction of active AMPKAR for parameter estimation; however, net AMPK activity is more physiologically relevant. To establish AMPK activity as a valid readout, we sought to verify how the AMPKAR sensor response correlates with AMPK kinase activity. For Model 6, we defined the net AMPK activity as

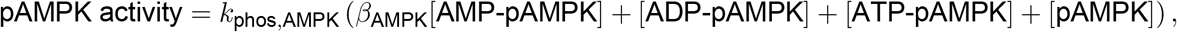

where the parameters *k*_phos,AMPK_ and *β*_AMPK_ were sampled from the posterior distribution. First, we found that the strength of ATP hydrolysis and calcium stimuli led to proportional increases in AMPK activity and decreases in the time-to-half max (Figure 5A–D). For a range of energy stress levels and calcium concentrations, the net AMPK kinase activity appeared to be more sensitive to changes in stimulus strength, did not saturate over relevant stimuli, and had faster kinetics than the AMPKAR sensor. Additionally, the response strength and timing were proportional for the AMPKAR and AMPK activity responses over moderate stimuli (Supplement Figure 5E-F); however, the sensor response began to saturate at higher input levels, whereas AMPK activity did not.

**Figure 5:**
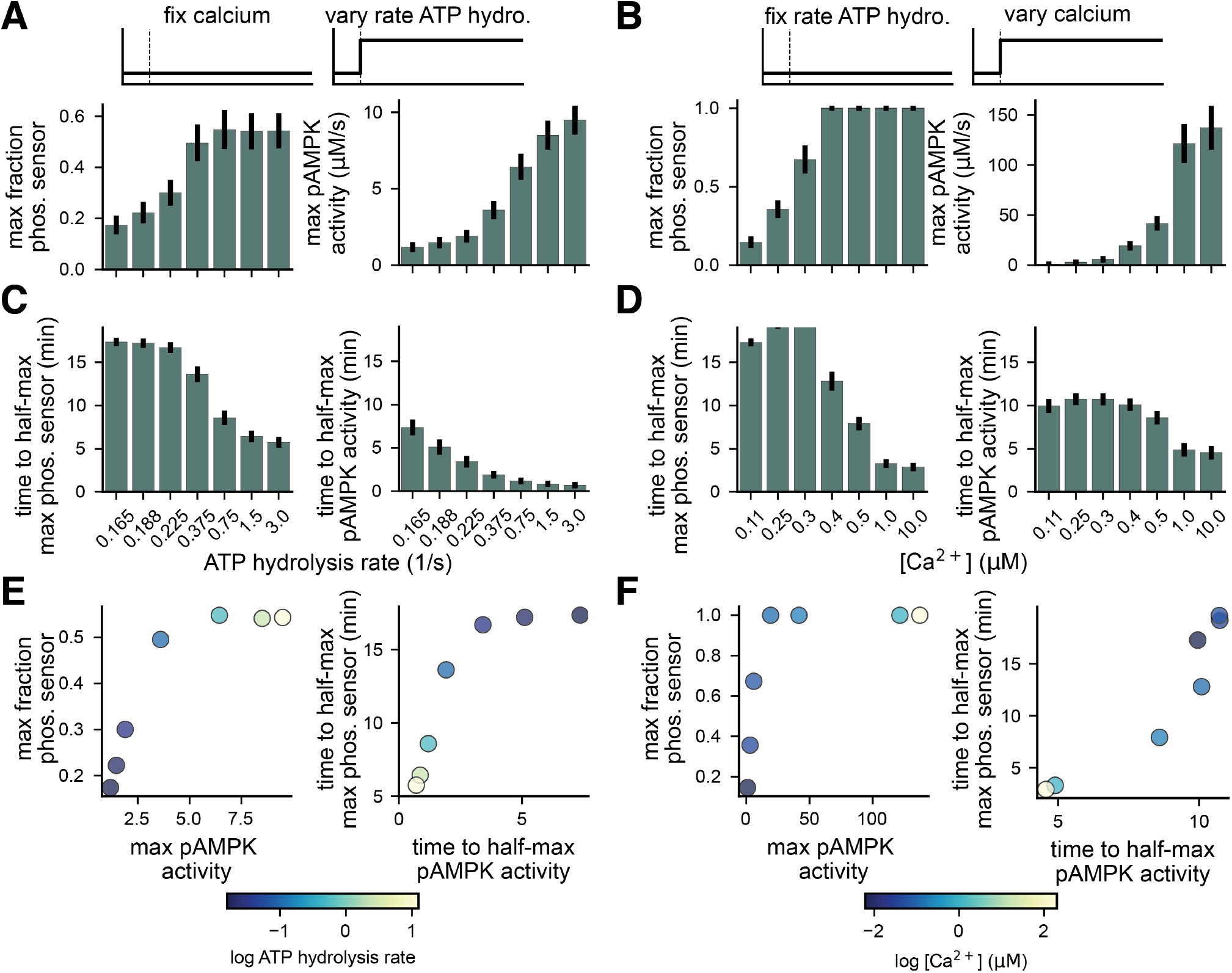
AMPKAR sensor and pAMPK activity vary proportionally with input strength. (**A**-**B**) Maximum fraction of activated AMPKAR sensor (left) and maximum pAMPK activity (right) as a function of ATP hydrolysis rate with fixed calcium (A) and as a function of calcium concentration with fixed ATP hydrolysis rate (B). (**C**-**D**) Time to half-maximum fraction of activated AMPKAR sensor (left) and maximum pAMPK activity (right) as a function of ATP hydrolysis rate with fixed calcium (C) and as a function of calcium concentration with fixed ATP hydrolysis rate (D). (**E**-**F**) Direct comparison of maximum values (left) and time to half-maximum values (right) as a function of ATP hydrolysis rate (E) and calcium concentration (F).

To understand how AMPK activity responds to more physiologically-relevant stimuli than 2-DG, we simulated transient increases in ATP hydrolysis and intracellular calcium. To do so, we applied a five-second pulse of increased intracellular calcium and ATP hydrolysis and let the system evolve for a total of 40 seconds (Figure 6A). We found that the area under the curve of pAMPK activity over the entire 40-second simulation was proportional to the calcium level, but not to the ATP hydrolysis level (Figure 6B). Interestingly, there appeared to be a threshold level of increased ATP hydrolysis between 2.0 and 5.0 mM/s that is necessary for the stimulus to begin to increase AMPK activity. We hypothesized that this discrepancy might be caused by slower dynamics in the metabolism module than in calcium-driven AMPK activation.

**Figure 6:**
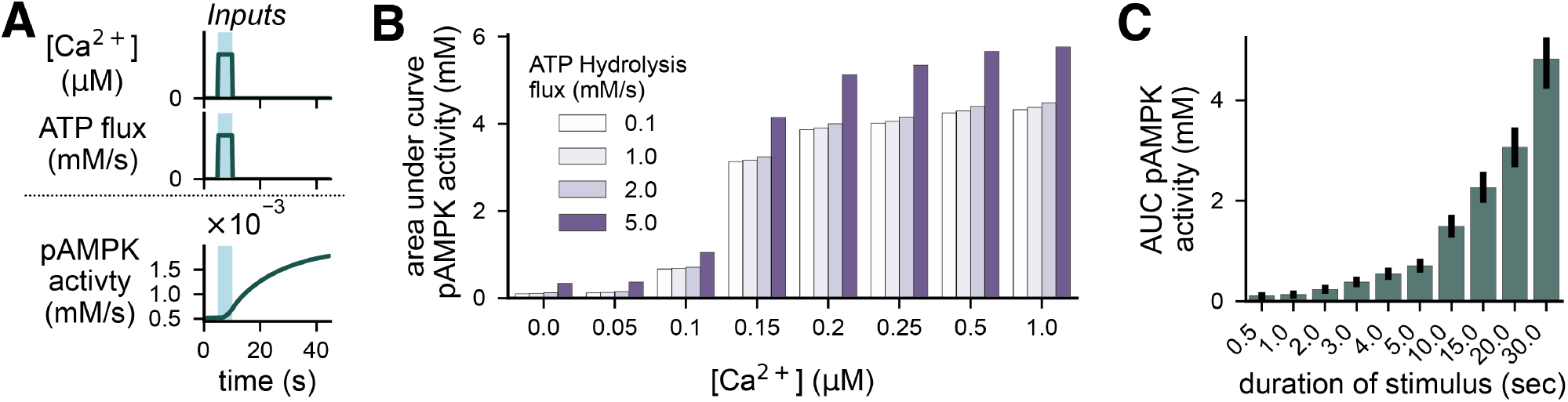
Stimulus duration induces greater changes in AMPK activity than stimulus strength. (**A**) Example of 10-second pulses of increased intracellular calcium and ATP flux and the corresponding trajectory of AMPK activity in the following 40 seconds. Blue shading indicates the time during which the stimulus is active. (**B**) Total AMPK kinase activity over 10 minutes as measured by the area under the curve of active pAMPK and AMP-pAMPK multiplied by the corresponding rate constants as a function of calcium and ATP flux stimuli. (**C**) Area under the curve of AMPK activity over 10 minutes as a function of stimulation duration for a fixed stimulus strength with [Ca^2+^] = 0.1 µM and ATP flux = 0.1 mM/s.

Next, the discrepancy between the effect of ATP hydrolysis and calcium led us to explore how the duration of a transient input affects AMPK activity. To do so, we fixed the strength of ATP hydrolysis input to 2.0 mM/s and the calcium input to 0.1 µM and varied the input duration from 0.5 to 30 seconds. We found that increasing the duration of the input led to proportional increases in AMPK activity, which were only obtained with calcium inputs at higher concentrations with a 5-second stimulus (Figure 6C). Additionally, we repeated these simulations, with either ATP hydrolysis or calcium fixed at baseline, to determine whether energy stress or calcium is more important during longer-duration stimuli. We found that calcium elevations appear to be responsible for most of the AMPK activity at short and moderate durations (0.5–10.0 seconds; Supplemental FigureS8). However, at longer durations, total AMPK activity with simultaneous stimuli appeared to be greater than the sum of the individual stimuli. This effect is likely because an increase in AMP concentration due to ATP depletion increases the effectiveness of calcium-induced CaMKK2 activity (Supplemental Figure S6) and thus increases AMPK activity.

Finally, we investigated how AMPK responds to periodic inputs, which are inspired by physiological signals in the myocyte during exercise [84]. To do so, we fixed the strength of ATP hydrolysis input to either 0.1 mM/s (low) or 1.0 mM/s (high) and the calcium input to 0.1 µM (low) or 0.25 µM (high) and varied the stimulus period over two timescales. On the shorter timescale, with higher frequencies, we applied a five-millisecond pulse and varied the period between the pulses from 0.006–0.2 seconds for a total of 10 seconds (Figure 7A). On the longer timescale, with lower frequencies, we applied a 5-second pulse for a total of 10 minutes (Figure 7D). For both timescales, we found that total AMPK activity measured by the area under the curve increases proportionally, but nonlinearly, with the input frequency (Figure 7B,D). However, at the high stimulus level, the total AMPK activity began to saturate for the short timescale inputs, but not for the long timescale inputs. These results suggest that AMPK responds proportionally to the input frequency.

**Figure 7:**
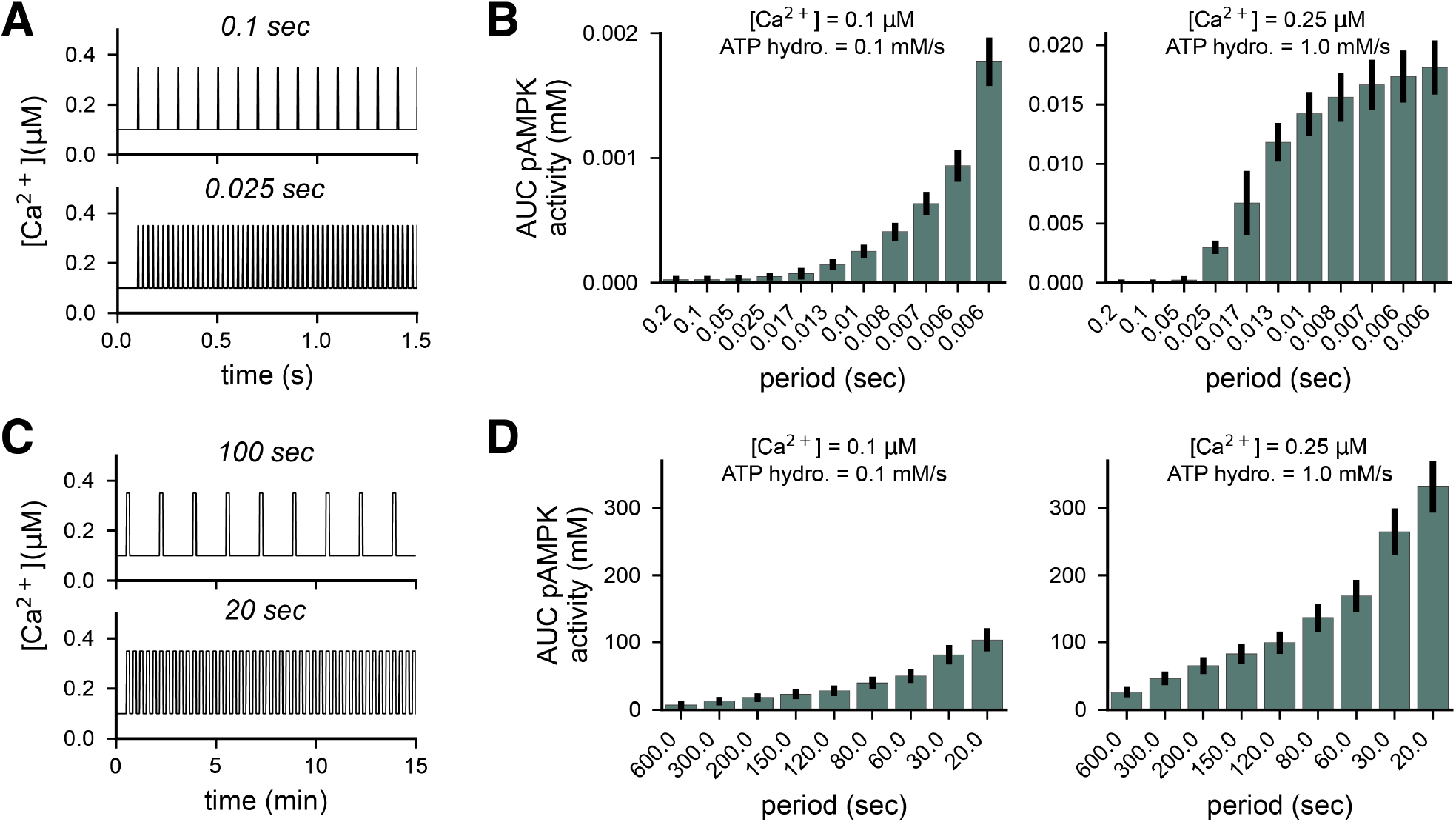
AMPK activity varies as a function of input frequency. *Short-timescale:* (**A**) Example of the first 2.5 seconds of a 10-second-long 10 Hz (top) and 40 Hz (bottom) Ca^2+^ stimuli. All pulses are 5 ms long. (**B**) AUC of AMPK activity over the 10 seconds as a function of stimulus frequency for a low-strength stimulus (left) and a high-strength stimulus (right). *Long-timescale:* (**A**) Example of the first 15 minutes of a 30-minute-long 0.6 min^−1^ (top) and 3 min^−1^ (bottom) Ca^2+^ stimuli. All pulses are 5 s long. (**B**) AUC of AMPK activity over the 30 minutes as a function of stimulus frequency for a low-strength stimulus (left) and a high-strength stimulus (right). For all panels, the low-strength stimulus is [Ca^2+^] = 0.1 µM and ATP flux = 0.1 mM/s, and the high-strength stimulus is [Ca^2+^] = 0.25 µM and ATP flux = 1.0 mM/s.

### 3.4 AMPK acts as an input signal integrator during short-timescale exercise

Finally, to demonstrate the applicability of our calibrated AMPK signaling model, we used the model to predict myocyte AMPK activity during exercise. Although AMPK is activated by a combination of energy stress and intracellular Ca^2+^, the role of AMPK in regulating metabolism in the muscle remains contested, with recent evidence suggesting that AMPK does not modulate glucose uptake and lipid metabolism during exercise [48, 49]. However, AMPK is responsible for post-exercise metabolic adaptation [25, 50–52]. Here, we investigated how AMPK would be activated by physiologically relevant energy stresses and Ca^2+^ inputs during simulated exercise.

Muscle activation is an inherently multiscale process that involves coordination between muscle fibers, myocytes, and motor units [85]. At the cellular level, muscle activation is driven by rapid, high-frequency neurological inputs that induce rapid changes in intracellular Ca^2+^ to drive force generation and, in turn, rapid changes in ATP consumption [84]. On a macroscopic scale, muscle contraction requires lower-frequency, slower bouts of increased myocyte activity, such as during a running stride or one repetition of a resistance exercise. Therefore, exercise induces changes in intracellular calcium that evolve over multiple timescales and can include high-frequency information of myocyte activation and lower-frequency information of muscle contraction. Different exercise conditions induce different myocyte activations at varying frequencies. For example, resistance training ranges from 10 to 50 Hz, and high-intensity interval training ranges from 60 to 180 Hz [86].

First, we simulated how AMPK responds to rapid changes in calcium and ATP hydrolysis during short-time-scale muscle activations. To do so, we simulated muscle activation during a resistance-like (Figure 8A) and a running-like (Figure 8B) stimulus using a detailed model of exercise-driven calcium signaling developed by Francis et al [87]. Briefly, this multicompartmental model captures action-potential-driven calcium dynamics and includes store-operated calcium release and cross-bridge cycling. Applying the predicted calcium and ATP hydrolysis transient to our AMPK signaling model allowed us to determine how AMPK responds to physiological inputs. The resistance stimulus consisted of 10 sets of 3 seconds of 40 Hz activation, followed by 3 seconds of rest, for a total of 60 seconds. The running stimulus was 0.1625 seconds of 100 Hz activation for each 0.65-second stride, repeated for a total of 60 seconds. We predicted AMPK activity in response to these exercise stimuli by simulating our model using the predicted calcium and ATP hydrolysis rates as inputs.

**Figure 8:**
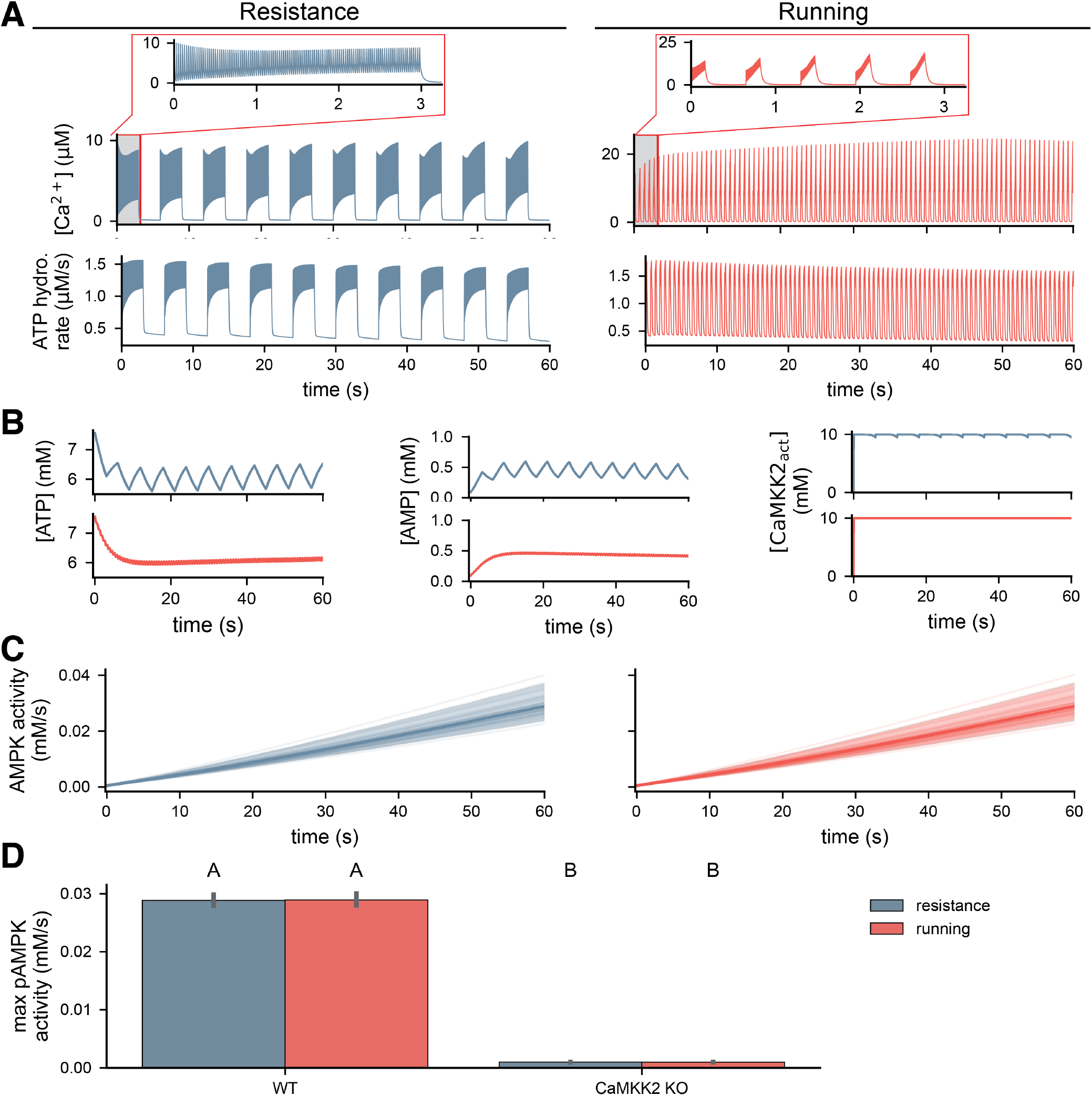
AMPK is robust to frequency differences during resistance and running exercise stimuli during short time-scale muscle activation. (**A**) Trajectories of [Ca]^2+^ and ATP hydrolysis rate during simulated resistance (left) and running (right) like stimuli. Simulations from myocyte exercise model from [87]. The inlay shows the first three seconds of the calcium. (**B**) Predicted ATP, AMP and activated CaMKK2 concentrations for each exercise type. (**C**) Posterior densities of predicted pAMPK activity during exercise stimuli. (**D**) Maximum pAMPK activity during exercise for wild-type and CaMKK2 KO conditions. Letters indicate statistical significance; groups with different letters are significantly different, and groups with like letters are significantly different. Significance was evaluated with a pairwise t-test and assessed at the level of 0.05.

Interestingly, the entire AMPK signaling pathway acted as a low-pass filter on the input signals. Upstream AMP, ATP, and active CaMKK2 only responded to slower variations in calcium and ATP hydrolysis induced by repetitions of contraction (resistance) or stride (running; Figure 8B). This aligns with previous findings that ligand-activated kinases, such as CaMKK2, can act as integrators and, thus, low-pass filter input signals [88]. In response, the predicted AMPK activity was nearly identical for the two exercise conditions (Figure 8C). Interestingly, the predicted AMPK activity trajectory only showed minor oscillations during the resistance condition. Quantitatively, the maximum AMPK activity was not significantly different in both conditions (WT; Figure 8C). Additionally, we found that exercise-induced AMPK activity depended strongly on CaMKK2, with CaMKK2 knockout abolishing nearly all AMPK activity (CaMKK2 KO; Figure 8C). These results suggest that they respond to exercise-related inputs in a manner that is robust to the high-frequency information in the signal.

### 3.5 AMPK activity is exercise-type dependent during long-timescale exercise

Finally, we investigated how AMPK responds to exercise-related stimuli at longer timescales. Sprint, endurance, and resistance exercise on the minute-to-hours scale have been shown to induce varied levels of AMPK activation both in vivo [25] and computationally [89]. In these studies, endurance induced greater AMPK activity than resistance, which was greater than that induced by sprinting. Here, we aimed to investigate AMPK activity in response to similar stimuli using our model. To do so, we simulated the three exercise conditions (Figure 9A), which mimic the timing of those prescribed in [25] and have levels informed by the average values of the stimuli in Figure 8A. The sprint condition was three 30-second intervals with a Ca^2+^ concentration of 10 µM and a rate of ATP hydrolysis of 3 1/s. The resistance condition was six 1-minute intervals with a Ca^2+^ concentration of 3 µM and a rate of ATP hydrolysis of 1 1/s. The endurance condition was one 90-minute interval with a Ca^2+^ concentration of 0.5 µM and a rate of ATP hydrolysis of 0.25 1/s.

**Figure 9:**
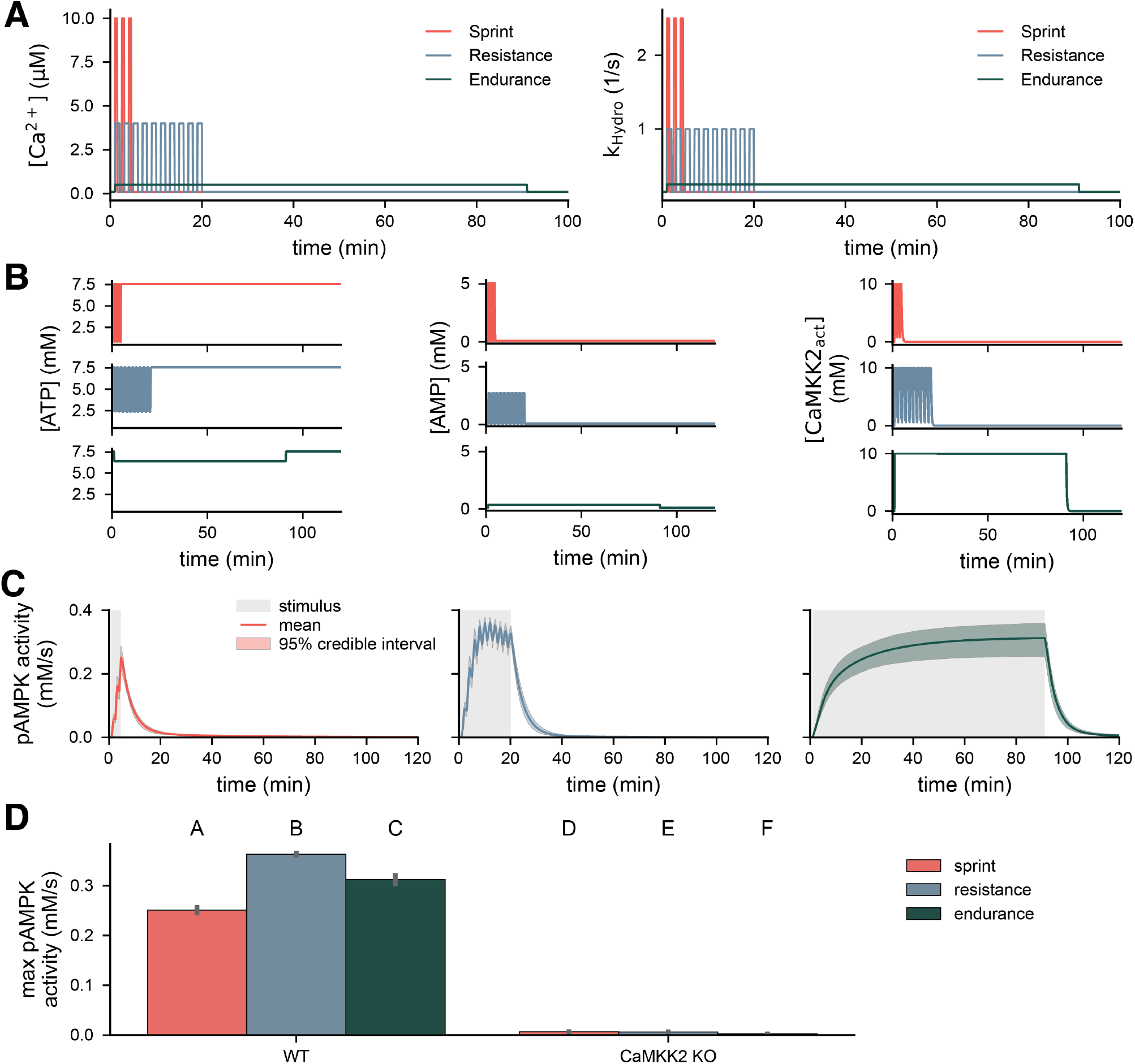
AMPK activation during long time-scale exercise stimuli depends on stimulus frequency and duration. (**A**) Long-time scale exercise stimuli during sprint (3 × 30s), resistance (6 × 1min) and endurance (1 × 90min). (**B**) Predicted ATP, AMP, and activated CaMKK2 concentrations for each exercise type. (**C**) Posterior densities of predicted pAMPK activity during exercise stimuli. Maximum pAMPK activity during exercise for wild-type and CaMKK2 KO conditions. Letters indicate statistical significance; groups with different letters are significantly different, and groups with like letters are significantly different. Significance was evaluated with a pairwise t-test and assessed at the level of 0.05.

Interestingly, these longer-timescale exercise stimuli induced stimulus-dependent AMPK activity. All three stimuli were slow enough to induce oscillatory (sprint and resistance) or step-wise (endurance) responses in AMP, ATP, and active CaMKK2 (Figure 9A). The subsequent trajectories of AMPK activity appeared to vary as a function of stimulus duration. Sprint induced a short and weaker response, resistance induced a stronger response, and endurance induced a moderate response. In all three trajectories, AMPK activity decreased to baseline after the stimulus duration. Resistance led to the highest maximum AMPK activity (WT; Figure 9A). Interestingly, endurance had higher maximum activity than sprint, even though the strength of the sprint stimulus was more than 10 times larger than the endurance stimulus. We additionally observed that AMPK activity depended strongly on CaMKK2 (CaMKK2 KO; Figure 9A). This result further suggests that AMPK integrates upstream signals and responds proportionally to the duration of the stimulus.

## Discussion

In this work, we developed a new mechanistic model of AMPK activation during energy stress. We took a data-informed approach to model development and applied Bayesian uncertainty quantification (Figure 2) to leverage recent high-quality experimental AMPK activity data [47]. First, we found that the choice of kinetic formulation used in a model impacted parameter identifiability and that there is no clear *best* formulation from that viewpoint. Next, we developed six related AMPK signaling models and found that the model using Michaelis-Menten kinetics and assuming that AMP and ADP act through nonessential mechanisms to modulate AMPK activity was most consistent with the data. Using this *best fit* model, we found that AMPK was sensitive to the timing of input stimuli, especially for energy-stress inducing inputs. In the context of exercise, we found that AMPK was robust to high-frequency variations in the input signal over short timescales but showed exercise dependence over longer timescales.

Uncertainty quantification, including identifiability analysis, prior elicitation, Bayesian parameter estimation, and Bayesian model selection, played a key role in developing the new model of AMPK signaling. In particular, UQ methods allowed us to utilize available experimental data to select a model and constrain its predictions. While previous applications of UQ in systems biology have enabled parameter estimation [40, 90, 91] and model selection [28, 55, 92], this work is unique in that we apply an end-to-end UQ workflow during the entire model development process. A key advancement here is the use of predictive prior elicitation and prior predictive sampling to iteratively refine the prior densities and values of the nonidentifiable parameters (Supplemental Section S1.3). This additional step addressed two challenges of our previously developed Bayesian parameter estimation workflow. First, predictive prior elicitation provided a structured approach to construct semi-informative priors when very little information is known about model parameters *a priori*. Second, iterative refinement of fixed nominal values allowed us to update nominal values for parameters that are mathematically nonidentifiable but still influence model predictions. While the UQ workflow presented here provided an end-to-end approach for data-informed model development, we did not *close the loop* and use the findings to update the model structures. To that end, all of the predictions were only as good as the set of input models. Methods for data-driven modeling, such as model discovery [93], hybrid mechanistic neural network models [94], and continuous model exploration with jump MCMC [95], can enable model development directly from data.

The mechanistic AMPK models capture AMPK activation by upstream kinases (LKB1 and CaMKK2) and activity control by adenine nucleotides. We chose to limit the model’s scope to the core of the AMPK signaling pathway to investigate the mechanisms by which adenine nucleotides control AMPK. However, as a hub of metabolic signaling, AMPK acts on several targets that can induce feedback to modulate AMPK activity, especially during longer timescale signaling events. For example, feedback through ULK1 and mTOR signaling could lead to net changes in AMPK activation [9, 96]. Furthermore, through its action on STIM1, AMPK could modulate Ca^2+^ signaling, which could, in turn, affect calcium-driven AMPK activation [97]. Future modeling efforts should focus on embedding mechanistic AMPK signaling models into the larger metabolic signaling system, which could be achieved, for example, by replacing the phenomenological representation with our mechanistic model [9–11]. Additionally, the role of AMPK activation in modulating energy uptake during exercise remains debated [49], so it is important to consider our predictions of exercise-induced AMPK activity fit into the broader myocyte metabolic signaling network. To better understand this, further modeling efforts should incorporate the relevant mechanisms and data necessary to investigate the roles of AMPK during exercise.

As experimental capabilities continue to improve, there is an ever-growing need to develop well-constrained models to integrate and interpret available data. Here, we showed how uncertainty quantification enables data-informed model development for signaling pathways where few previously developed models had been published. We believe that this work serves as a starting point for future modeling and development of new UQ methods in systems biology. As we showed here, the application of UQ in systems biology has the potential to improve modeling and better constrain model-based predictions to experimental data ans drive new discoveries in intracellular signaling.

## Code Availability

We provide all the code and instructions to reproduce these results in the following GitHub repository https://github.com/natejlinden/AMPK.

## Acknowledgments

The authors would like to give special thanks to Drs. Danielle Schmitt (University of California, Los Angeles) and Allen Leung for many initial discussions in this area of research. Additionally, they would like to thank Dr. Emmet Francis, Juliette Hamid, and Anne Lyons for their insights and helpful discussions. NLS acknowledges support from the National Institute of Biomedical Imaging and Bio-engineering (NIBIB) of the National Institutes of Health (NIH; https://www.nibib.nih.gov) under award number T32EB9380 and a UCSD Sloan Scholar Fellowship from the Alfred P. Sloan Foundation (https://sloan.org). NLS and PR acknowledge support Wu Tsai Human Performance Alliance at UCSD.

## S1 Supplemental Text

### S1.1 Two-reaction models

To further investigate how the choice of kinetic formulation affects parameter identifiability, we examined models of a two-reaction loop that was inspired by a phosphorylation-dephosphorylation loop. The models represented the reactions

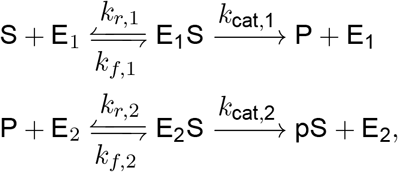

where S is the substrate, E_1_ and E_2_ are the enzymes, and P is the product. The product of the first reaction is the substrate of the second P and the product of the second reaction is the original substrate S. To investigate the effects of the kinetic formulation, we constructed nine sets of models that represented the two-reaction systems. The models differed in the kinetic formulation used to represent either reaction, specifically mass action, Michaelis-Menten, or Hill-type. For example, one of the models employed mass action kinetics for both reactions, while another used mass action for the first reaction and the Michaelis-Menten equation for the second. Supplemental Table 3 summarizes the models and the identifiability results.

### S1.2 AMPK model details

In this section, we provide additional rationale and derivations for the six AMPK signaling models. Unless otherwise noted, all relevant assumptions are outlined in the manuscript.

#### S1.2.1 Module 1: Cellular metabolism and energy stress

We adapted the simplified metabolism models from Coccimiglio et al. [30] and Lueng et al. [9]. Our metabolism model includes phenomenological representations of oxidative phosphorylation, glycolysis, ATP hydrolysis, adenylate kinase, and creatine kinase. First, we represented ATP generation by oxidative phosphorylation with a first-order irreversible reaction from ADP to ATP, which we modeled with the Hill-type equation originally derived in [98]. Second, we represented glycolysis with an irreversible first-order reaction from ADP to ATP, which we modeled with mass action kinetics. Third, we represented net ATP consumption due to hydrolysis with a single hydrolysis reaction from ATP to ADP, which we modeled with mass action kinetics. Fourth, we included both the adenylate and the creatine kinase reactions to ensure the correct buffering of adenine ratios [99]. The adenylate kinase reaction is

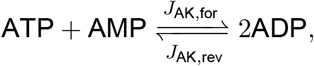

which we modeled with a bi-bi reaction mechanism originally derived in [99]. Finally, the reversible creatine kinase reaction is

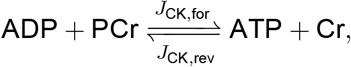

which we modeled with a Cleland bi-bi reaction that was originally derived in [99]. Table 4 lists the fluxes, *J*_OhPhos_, *J*_Gly_, etc., for these reactions.

The associated differential equations for ATP, ADP, AMP, and PCr, are defined as,

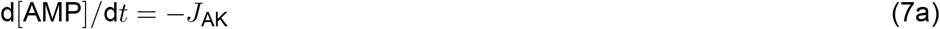

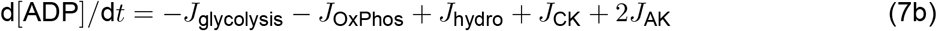

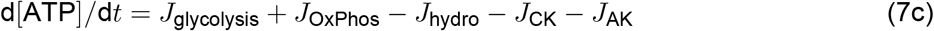

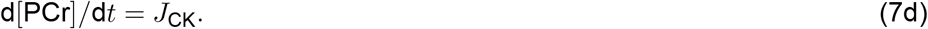

Throughout this work, we assume that the initial concentrations of AMP, ADP, and ATP are 7.03 mM, 1.11 mM, and 7.9 × 10^−2^ mM, respectively. Supplemental Figure S1A shows transient AMP, ADP, and ATP concentrations from an arbitrary initial condition to the steady state taken as the initial condition in this work. The cellular metabolism model is the same across all models that we develop. Table 5 provides values for all of the Module 1 parameters. All parameter values were taken from sources, except for the rate of ATP production by glycolysis. We derived the glycolysis rate *k*_glycolysis_ roughly on experimental measurements of glycolytic flux from Salti et al. [100] Figure 2. In MEF cells, those authors measured a flux of 9.16 pmol/ATP/10^3^ cells, which we converted to mM/s by assuming a cellular volume of 4000 *µ*m^3^, which gives approximately 22 mM/s. We divided by twice initial concentration of ATP (7.03 mM) and rounded down to yield *k*_gly_ =1.5 1/s.

#### S1.2.2 Module 2: Adenine nucleotide binding and AMPK phosphorylation

The models include adenine nucleotide binding to AMPK, phosphorylation of AMPK, and dephosphorylation of AMPK. In this section, we describe additional model assumptions and outline the nonessential and essential activation/inhibition mechanisms used in the models.

##### Adenine nucleotide binding

We assumed that adenine nucleotide binding kinetics do not vary depending on the phosphorylation or binding state of AMPK. For example, in the models, the reactions

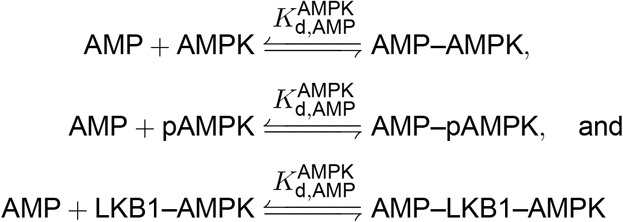

all occur with the same kinetics. We used the binding coefficients from [16] as initial values when constructing prior densities for these parameters.

##### AMPK phosphorylation by upstream kinases LKB1 and CaMKK2

We assumed that AMPK phosphorylation by LKB1 and CaMKK2 occurs as a standard enzyme-mediated reaction [38]. For example, phosphorylation by LKB1 would be described as

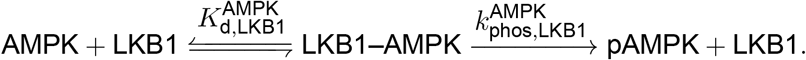

In order to capture the substrate-mediated effects of AXP binding, we assumed that AMP and ADP act as either essential activators (Models 1 and 2) or nonessential activators (Models 4–6) of upstream kinases. Essential activation assumes that the activator must be bound to the substrate for the reaction to proceed [38]. For example,

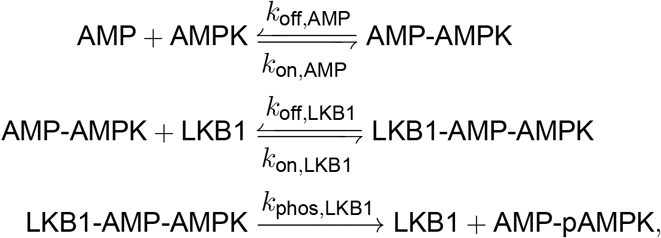

assumes that AMP is the essential activator of LKB1-induced phosphorylation. Alternatively, nonessential activation assumes that the activator increases the reaction rate or decreases the *K*_*d*_ between the enzyme and substrate [38]. For example, in the reaction

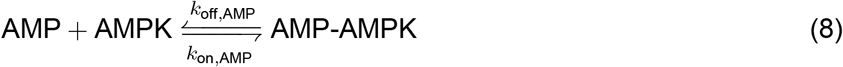

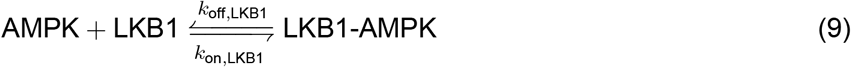

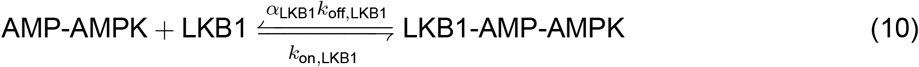

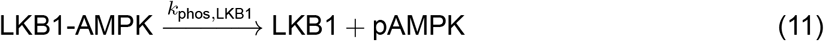

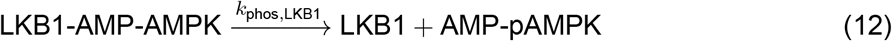

the activator AMP decreases the *K*_*d*_ by a factor of *α*_LKB1_ ≤ 1.

##### AMPK dephosphorylation

Similar to phosphorylation, we assumed that dephosphorylation of phosphorylated pAMPK occurs as a standard enzyme-mediated reaction. In Models 1 and 2, we assumed that AMP and ADP act as essential inhibitors of the dephosphorylation reaction, where AMP-pAMPK and ADP-pAMPK are not dephosphorylated. In Models 3–6, we assumed that AMP and ADP acts as nonessential inhibitors of the dephosphorylation reaction, either increasing the *K*_*d*_ (Models 3 and 4) or decreasing the reaction rate (Models 5 and 6).

### S1.3 Predictive prior elicitation and prior refinement

We encountered two key challenges in constructing prior densities for model parameters: (i) choosing priors that yield reasonable predictions when little prior information is available and (ii) assigning values to nonidentifiable parameters that we fix before estimation. Here, we took an approach that combined predictive prior elicitation with iterative updating of fixed parameters. Predictive prior elicitation is a qualitative approach that allows us to adjust the parameters of the prior densities such that the prior predictive distribution appears reasonable [58]. This allows us to refine our prior beliefs without utilizing any data. Briefly, the prior predictive distribution shows the distribution of data generated by the model given assumed prior densities [41]. In the remainder of this section, we outline the process used to refine prior densities and select values for non-identifiable parameters, using Model 2 as an example.

We began by using direct prior elicitation to choose default priors for uncertain parameters by specifying a range of plausible values (see Section 2) [58]. Using these priors for Model 2, we observed that the prior predictive density was unreasonable, because all of the predicted simulations were constantly zero (Supplemental Figure S2A). We hypothesized that the values of the nonidentifiable parameters, which we fixed to nominal values, could account for the observed behavior. To address this, we varied each of the five nonidentifiable parameters independently over several orders of magnitude and simulated the corresponding prior predictive densities. Supplemental Figure S2B shows the cumulative density functions of the final value of the predicted sensor response. We found that only *k*_AMPK_ affected the predictions, so we updated the nominal value for that parameter to 0.192 because that value yielded more reasonable predictions. We repeated this process for the remaining parameters and found that we only needed one additional iteration of updates, because only *k*_CaMKK_ affected predictions (Supplemental Figure S2C). Given that we were satisfied with the updated nominal values for nonidentifiable parameters, we next chose to refine the priors using predictive prior elicitation. Here, we simulated the prior density and iteratively updated the hyperparameters of the priors to yield more reasonable predictions. We refer the reader to the tutorial in the Preliz Python package, at the following link https://preliz.readthedocs.io/en/latest/examples/gallery/predictive_explorer.html for more details. Supplemental Figure S2D shows the resulting prior predictive distribution and Supplemental Figure S2E shows the densities of the final value in the simulation and the time to half max. We repeated this process for all six AMPK models (Supplemental Figure S3).

## S2 Supplemental Figures

**Figure S1:**
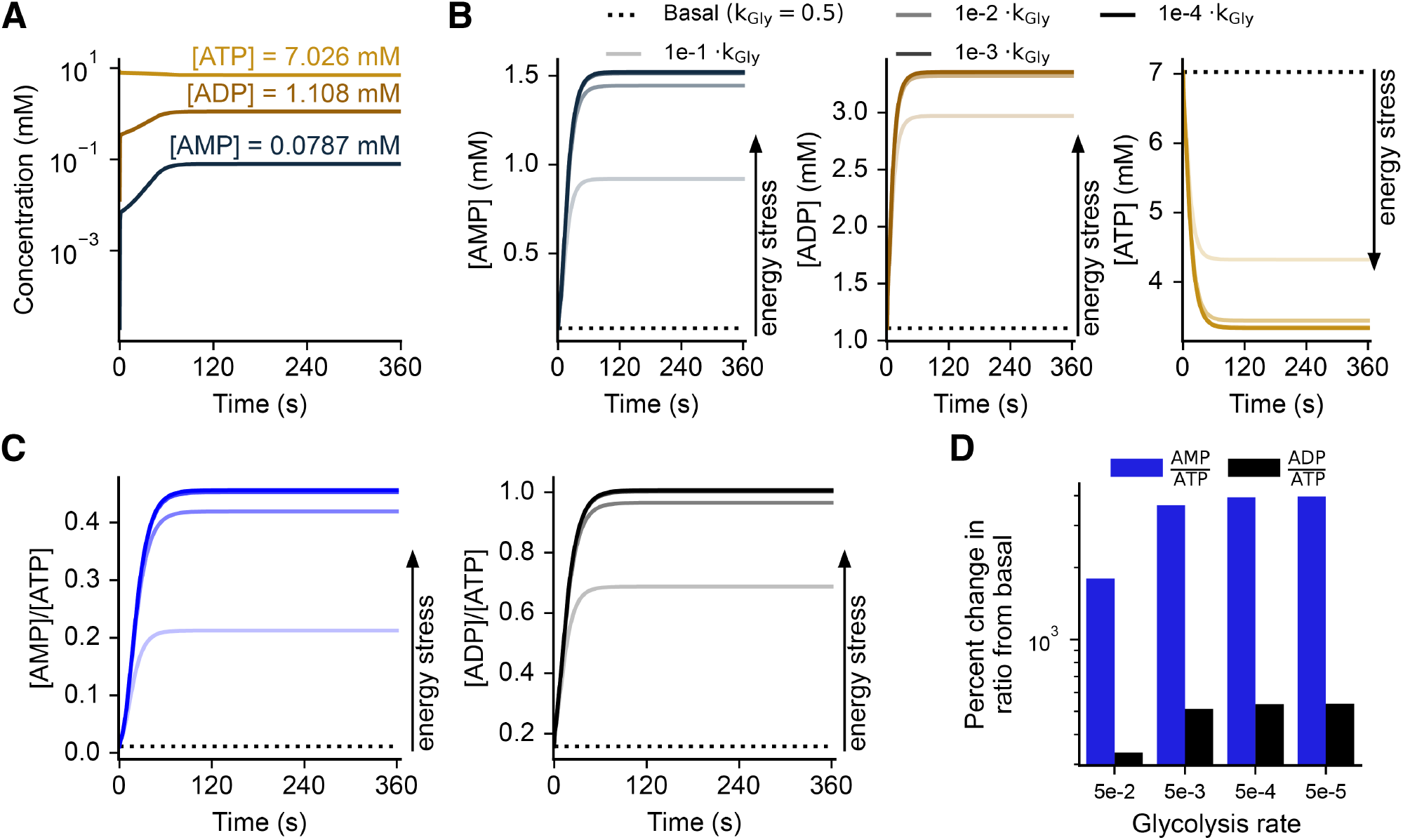
Simulated energy stress with decreased glycolytic ATP production increases the AMP/ATP and ADP/ATP ratios. (**A**) AMP, ADP, and ATP transients from arbitrary initial conditions to steady state values with basal ATP production. (**B**) Effect of energy stress due to decreasing the rate of glycolytic ATP production on AMP, ADP, and ATP concentrations over time. The rate of glycolysis is changed from the basal value at *t* = 0. (**C**) Effect of energy stress on transient AMP/ATP (blue) and ATP/ADP (black) ratios. Stimuli are the same as in B. (**D**) Percent change in AMP/ATP and ADP/ATP ratios from basal due to increased energy stress.

**Figure S2:**
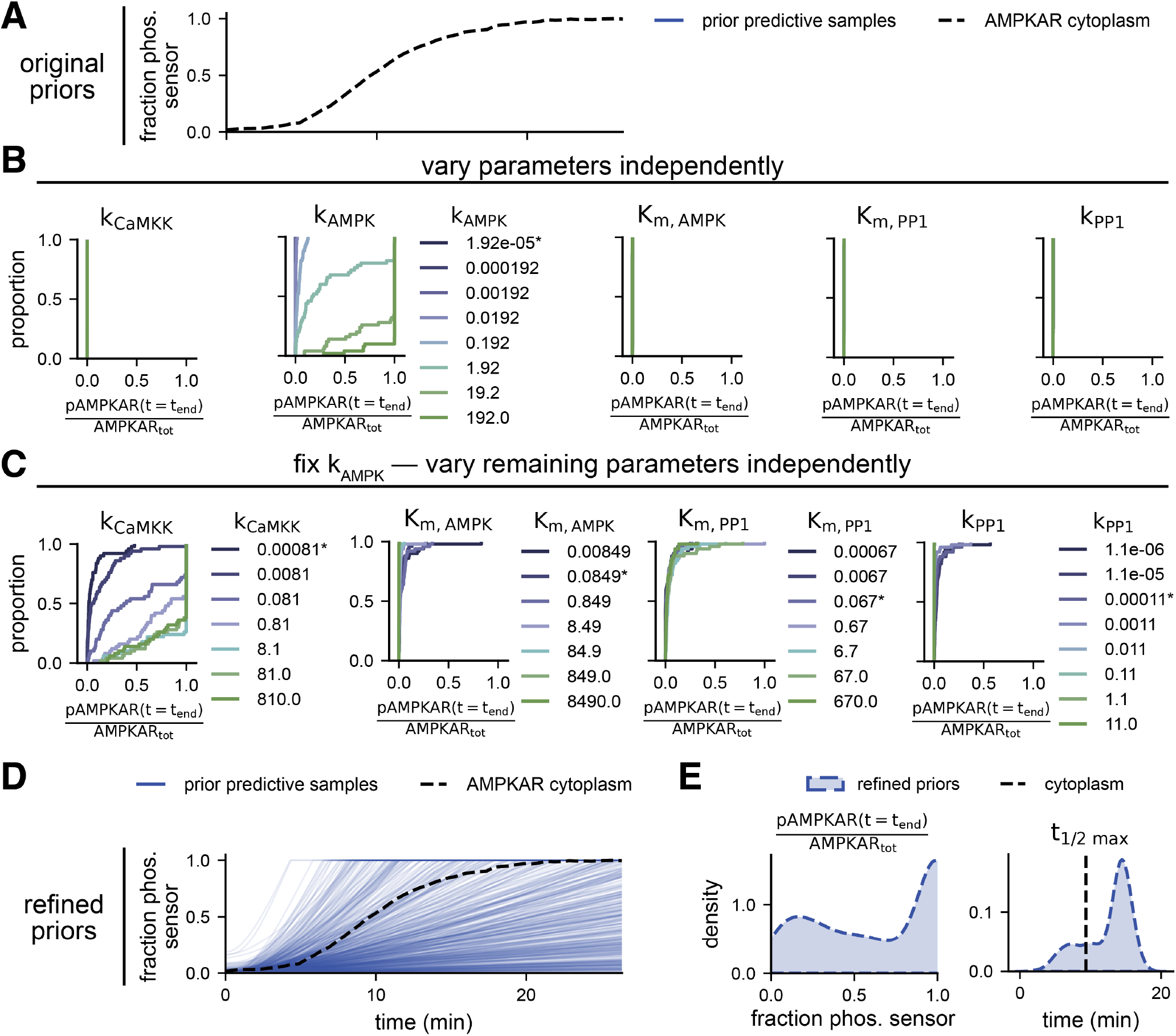
Prior refinement and nominal value selection for Model 2. (**A**) Prior predictive distribution of predictions for priors based-on parameter ranges before any refinement. All simulations are nearly zero for all time. (**B**) Empirical cumulative density functions (CDF) for the final fraction of activated AMPKAR from prior predictive samples with single-parameter variations of nonidentifiable parameters. Only *k*_AMPK_ affects the strength of AMPKAR response. (**C**) Empirical CDFs for the final fraction of activated AMPKAR from prior predictive samples with single-parameter variations of nonidentifiable parameters with *k*_AMPK_ fixed to 0.192. Only *k*_CaMKK_ affects the strength of AMPKAR response. (**D**) Prior predictive distribution of predictions for refined priors and updated values for *k*_AMPK_ and *k*_CaMKK_. (**E**) Estimated probability densities of key quantities after prior refinement.

**Figure S3:**
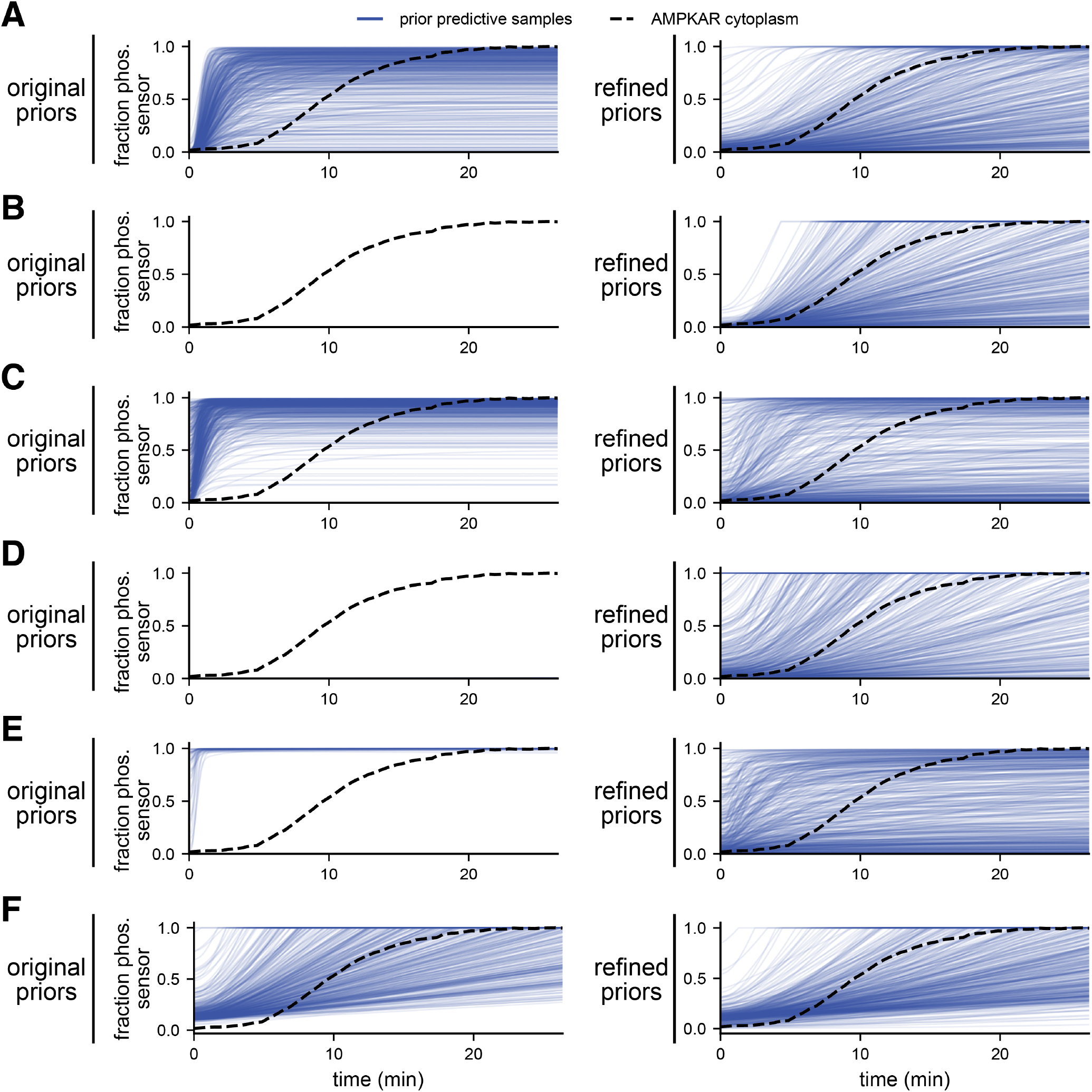
Prior predictive simulation samples with the original (left) and refined (right) priors for all models. Dashed black line shows the data mean. Transparent blue lines show simulations with 400 prior samples.

**Figure S4:**
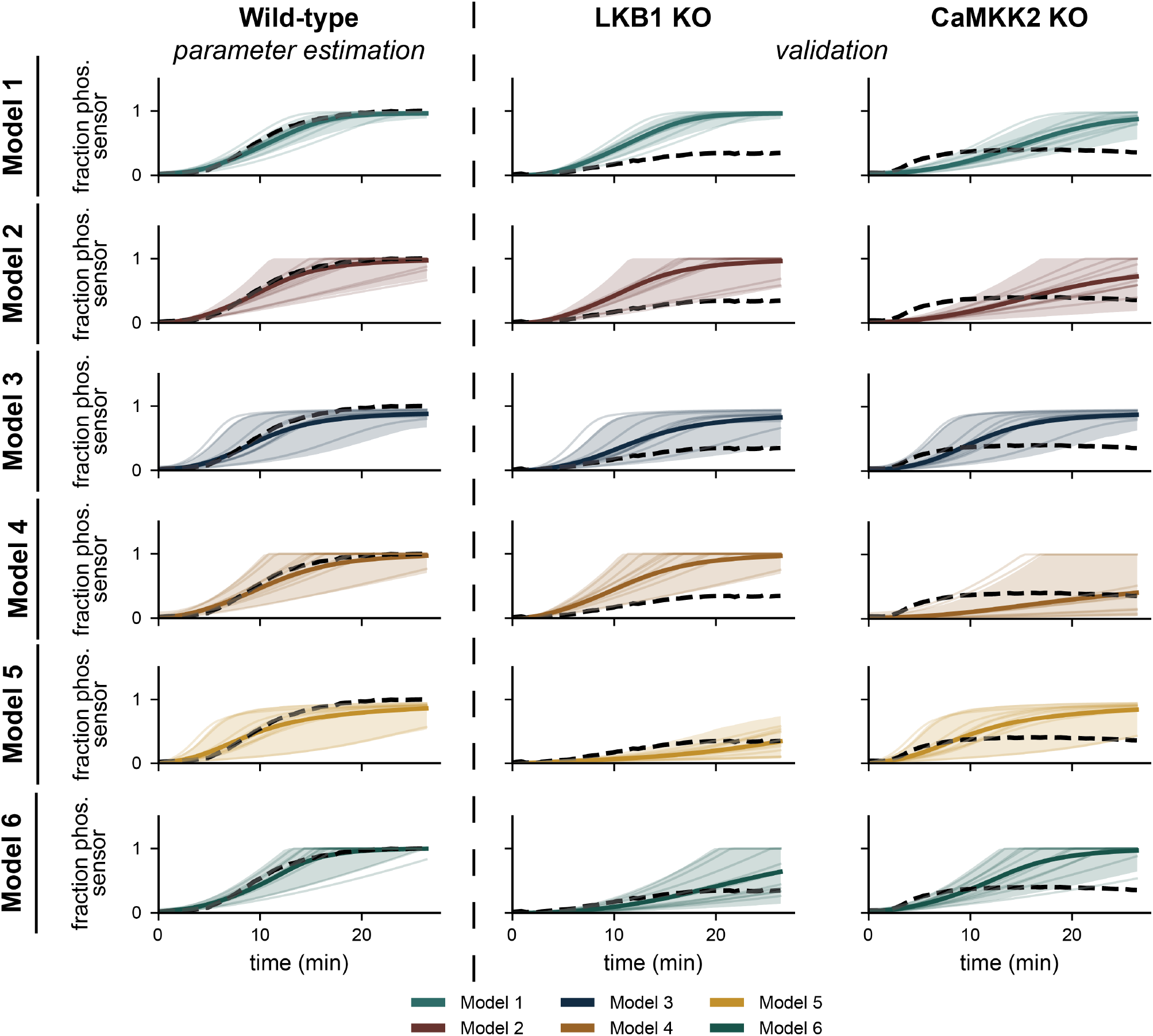
Posterior predictions for all models with parameters estimated with only wild-type data. Dashed black lines show the data mean. Solid colored lines show the posterior mean, the shaded band shows the 95% credible interval, and the transparent lines show 10 samples from the posterior density. LKB1 and CaMKK2 knockout conditions are forward predictions.

**Figure S5:**
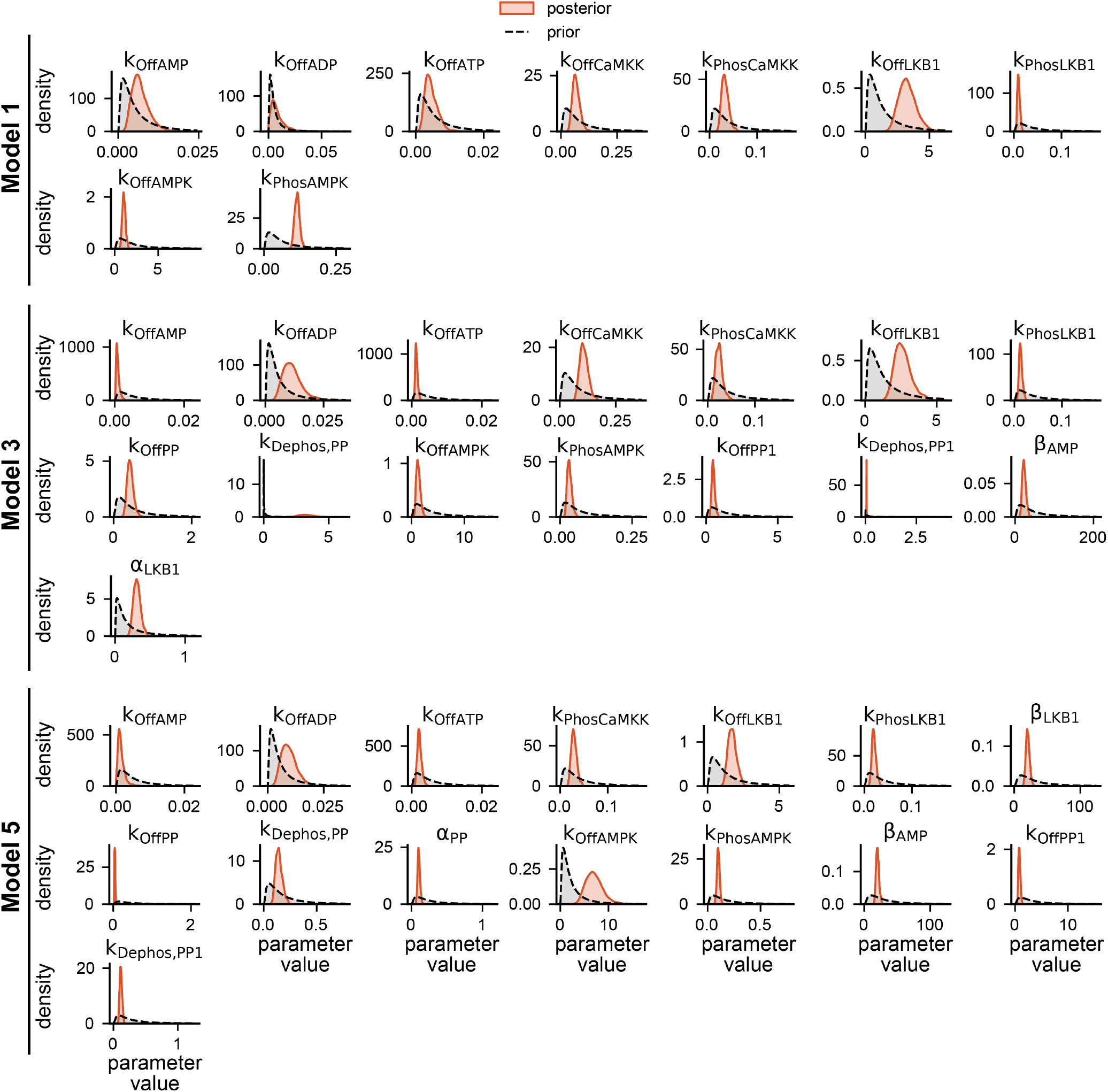
Estimated marginal posterior (orange) and prior (black dashed) densities of models with mass action kinetics, Models 1, 3, and 5. Densities were estimated by fitting a kernel density estimator to 1000 posterior samples. Posterior was sampled using the Pathfinder variational inference algorithm [63].

**Figure S6:**
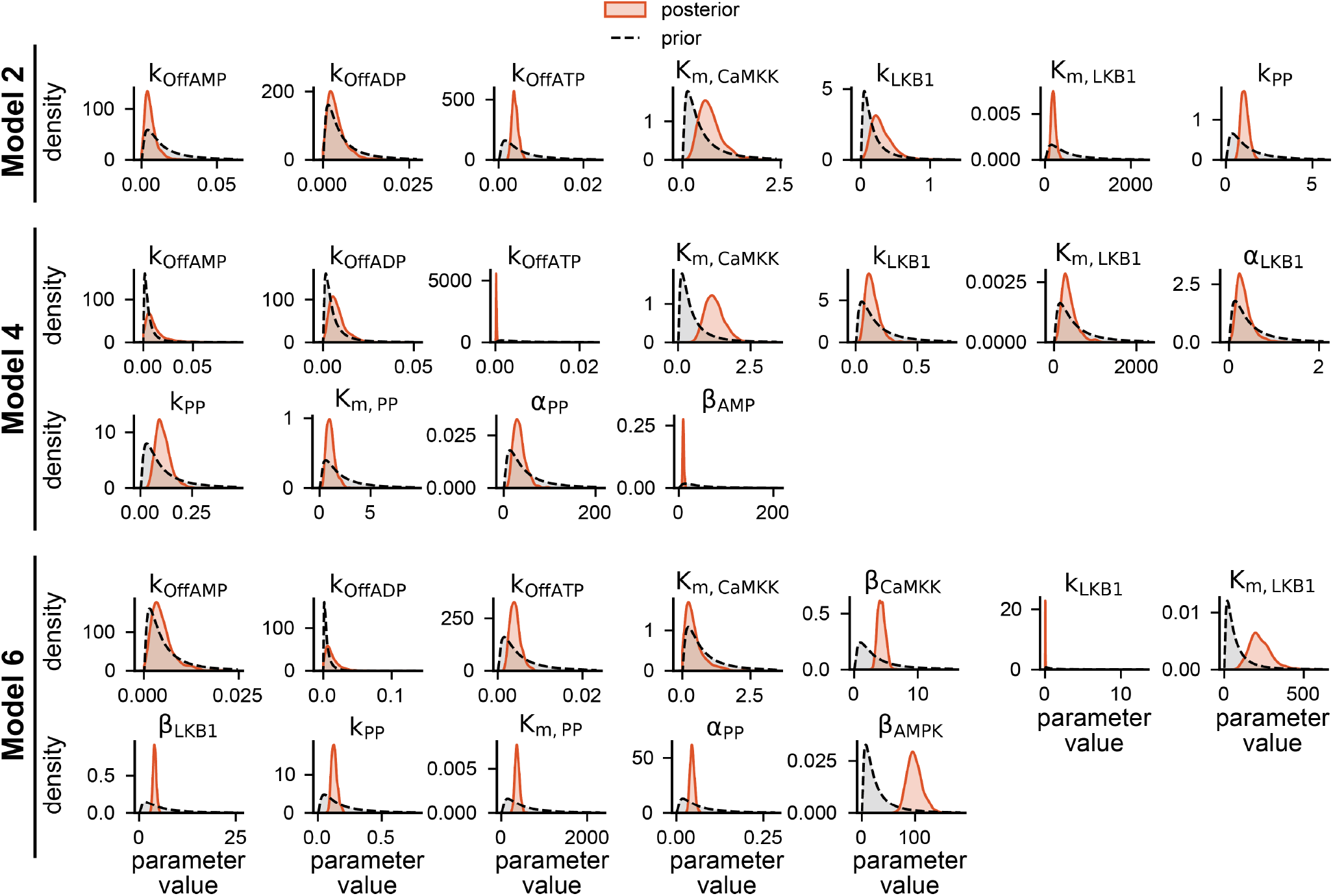
Estimated marginal posterior (orange) and prior (black dashed) densities of models with Michaelis-Menten kinetics, Models 2, 4, and 6. Densities were estimated by fitting a kernel density estimator to 1000 posterior samples. Posterior was sampled using the Pathfinder variational inference algorithm [63].

**Figure S7:**
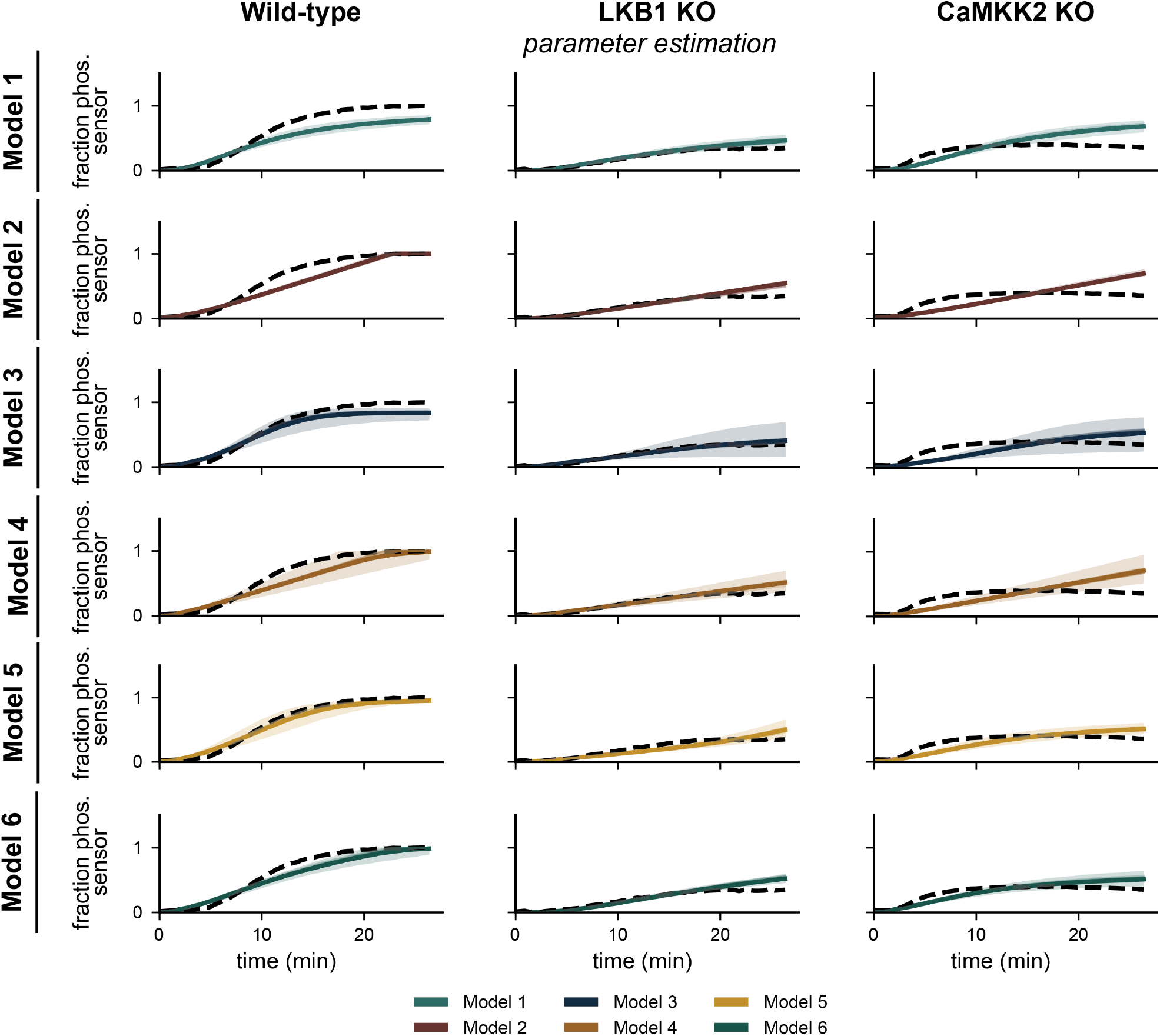
Posterior predictions for all models with parameters estimated with data from all three conditions simultaneously. Dashed black lines show the data mean. Solid colored lines show the posterior mean, the shaded band shows the 95% credible interval, and the transparent lines show 10 samples from the posterior density.

**Figure S8:**
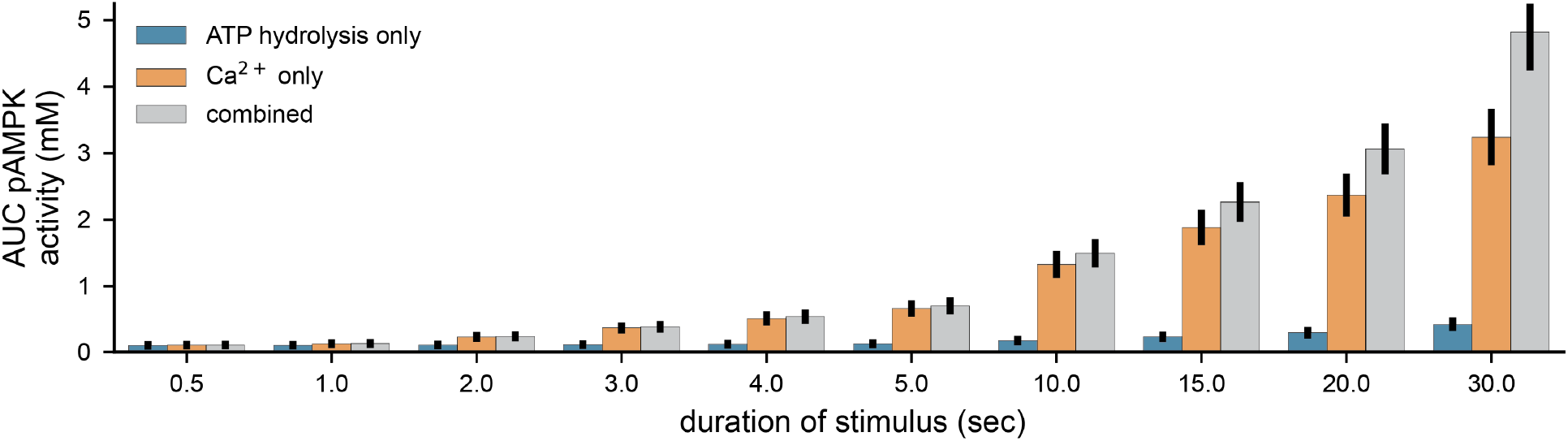
Area under the curve as a function of stimulus duration. Stimuli strengths were either [Ca^2+^] = 0.1 µM (Ca^2+^ only), ATP flux = 2.0 mM/s (ATP hydrolysis only), or both (combined).

## S3 Supplemental Tables

**Table 2:** Prior parameters for the single enzyme-mediated reaction models shown in Figure 3.

| Model | Parameter | Units | Prior |
| --- | --- | --- | --- |
| Mass action | $k_f$ | 1/(mM s) | Gamma ($\alpha = 3.14, \beta = 33.01$) |
| Mass action | $k_r$ | 1/s | Gamma ($\alpha = 3.14, \beta = 132.04$) |
| Mass action | $k_{cat}$ | 1/s | Gamma ($\alpha = 3.14, \beta = 33.01$) |
| Michaelis-Menten | $V_{max}$ | 1/s | Gamma ($\alpha = 4.45, \beta = 420.00$) |
| Michaelis-Menten | $K_m$ | mM | Gamma ($\alpha = 4.45, \beta = 1680.00$) |
| Hill-type | $V_{max}$ | 1/s | Gamma ($\alpha = 4.45, \beta = 4200.00$) |
| Hill-type | $K_m$ | mM | Gamma ($\alpha = 3.14, \beta = 132.04$) |
| Hill-type | $n$ | unitless | TruncatedNormal* ($\mu = 1.39, \sigma = 0.36, lb = 1, ub = \infty$) |

\* truncated normal distribution is truncated at one, so support is over $[1, \infty)$

**Table 3:** Identifiability of two-reaction models. Naming indicates the kinetic formulation used for the first model and the second reaction, respectively. For example, mass action-mass action uses mass action kinetics for both reactions. Abbreviations: MA–mass action, MM–Michaelis-Menten, Hill– Hill-type.

| Model | Locally identifiable<br>$y_{\text{obs}} = [P]$ | NOT identifiable<br>$y_{\text{obs}} = [P]$ | Locally identifiable<br>$y_{\text{obs}} = \frac{[P]}{[S]_{\text{tot}}}$ | NOT identifiable<br>$y_{\text{obs}} = \frac{[P]}{[S]_{\text{tot}}}$ |
| --- | --- | --- | --- | --- |
| MA–MA | $k_{f,1}, k_{r,1}, k_{\text{cat},1}, k_{f,2}, k_{r,2}, k_{\text{cat},2}$ | – | $k_{r,1}, k_{\text{cat},1}, k_{r,2}, k_{\text{cat},2}$ | $k_{f,1}, k_{f,2}$ |
| MA–MM | $k_{f,1}, k_{r,1}, k_{\text{cat},1}, V_{\text{max},2}, K_{m,2}$ | – | $k_{r,1}, k_{\text{cat},1}$ | $k_{f,1}, V_{\text{max},2}, K_{m,2}$ |
| MA–Hill | $k_{f,1}, k_{r,1}, k_{\text{cat},1}, V_{\text{max},2}, K_{m,2}$ | – | $k_{r,1}, k_{\text{cat},1}$ | $k_{f,1}, V_{\text{max},2}, K_{m,2}$ |
| MM–MM | $V_{\text{max},1}, K_{m,1}, V_{\text{max},2}, K_{m,2}$ | – | – | $V_{\text{max},1}, K_{m,1}, V_{\text{max},2}, K_{m,2}$ |
| MM–MA | $V_{\text{max},1}, K_{m,1}, k_{f,2}, k_{r,2}, k_{\text{cat},2}$ | – | $V_{\text{max},1}, K_{m,1}, k_{f,2}, k_{r,2}, k_{\text{cat},2}$ | – |
| MM–Hill | $V_{\text{max},1}, K_{m,1}, V_{\text{max},2}, K_{m,2}$ | – | – | $V_{\text{max},1}, K_{m,1}, V_{\text{max},2}, K_{m,2}$ |
| Hill–Hill | $V_{\text{max},1}, K_{m,1}, V_{\text{max},2}, K_{m,2}$ | – | – | $V_{\text{max},1}, K_{m,1}, V_{\text{max},2}, K_{m,2}$ |
| Hill–MA | $V_{\text{max},1}, K_{m,1}, k_{f,2}, k_{r,2}, k_{\text{cat},2}$ | – | $k_{r,2}, k_{\text{cat},2}$ | $V_{\text{max},1}, K_{m,1}, k_{f,2}$ |
| Hill–MM | $V_{\text{max},1}, K_{m,1}, V_{\text{max},2}, K_{m,2}$ | – | – | $V_{\text{max},1}, K_{m,1}, V_{\text{max},2}, K_{m,2}$ |

**Table 4:** Fluxes for the metabolism module (Module 1).

| Reaction | Flux | Reference |
| --- | --- | --- |
| Oxidative phos. | $J_{\text{OxPhos}} = \frac{V_{\text{max, OxPhos}} \left( \frac{[\text{ADP}]}{K_{\text{ADP}}} \right)^n}{1 + \left( \frac{[\text{ADP}]}{K_{\text{ADP}}} \right)^n}$ | [98] |
| Glycolysis | $J_{\text{glycolysis}} = k_{\text{glycolysis}} [\text{ADP}]$ | n/a |
| ATP Hydrolysis | $J_{\text{hydro}} = k_{\text{hydro}} [\text{ATP}]$ | [30] |
| Adenylate kinase<br>(forward) | $J_{\text{AK,for}} = \frac{\frac{V_{\text{AK,for}} [\text{ATP}] [\text{AMP}]}{k_{\text{mt}} \times k_{\text{mm}}}}{1 + \frac{[\text{ATP}]}{k_{\text{mt}}} + \frac{[\text{AMP}]}{k_{\text{mm}}} + \frac{[\text{ATP}] [\text{AMP}]}{k_{\text{mt}} \times k_{\text{mm}}} + \frac{2[\text{ADP}]}{k_{\text{md}}} + \frac{[\text{ADP}]^2}{k_{\text{md}}^2}}$ | [99] |
| Adenylate kinase<br>(reverse)* | $J_{\text{AK,rev}} = \frac{\frac{V_{\text{AK,rev}} [\text{ADP}]^2}{k_{\text{md}}^2}}{1 + \frac{[\text{ATP}]}{k_{\text{mt}}} + \frac{[\text{AMP}]}{k_{\text{mm}}} + \frac{[\text{ATP}] [\text{AMP}]}{k_{\text{mt}} \times k_{\text{mm}}} + \frac{2[\text{ADP}]}{k_{\text{md}}} + \frac{[\text{ADP}]^2}{k_{\text{md}}^2}}$ | [99] |
| Adenylate kinase | $J_{\text{AK}} = J_{\text{AK,for}} - J_{\text{AK,rev}}$ | [99] |
| Creatine kinase<br>(forward) | $J_{\text{CK,for}} = \frac{\frac{V_{\text{CK,for}} K_{iq} K_p [\text{ADP}] [\text{PCr}]}{K_{ia} K_b}}{1 + \frac{[\text{ADP}]}{K_{ia}} + \frac{[\text{PCr}]}{K_{ib}} + \frac{[\text{ATP}]}{K_{iq}} + \frac{[\text{ADP}] [\text{PCr}]}{K_{ia} K_b} + \frac{([\text{Cr}]_{\text{total}} - [\text{PCr}]) [\text{ATP}]}{K_{iq} K_p}}$ | [99] |
| Creatine kinase<br>(reverse)** | $J_{\text{CK,rev}} = \frac{\frac{V_{\text{CK,rev}} ([\text{Cr}]_{\text{total}} - [\text{PCr}]) [\text{ATP}]}{K_{iq} K_p}}{1 + \frac{[\text{ADP}]}{K_{ia}} + \frac{[\text{PCr}]}{K_{ib}} + \frac{[\text{ATP}]}{K_{iq}} + \frac{[\text{ADP}] [\text{PCr}]}{K_{ia} K_b} + \frac{([\text{Cr}]_{\text{total}} - [\text{PCr}]) [\text{ATP}]}{K_{iq} K_p}}$ | [99] |
| Creatine kinase | $J_{\text{CK}} = J_{\text{CK,rev}} - J_{\text{CK,for}}$ | [99] |

**Table 5:**
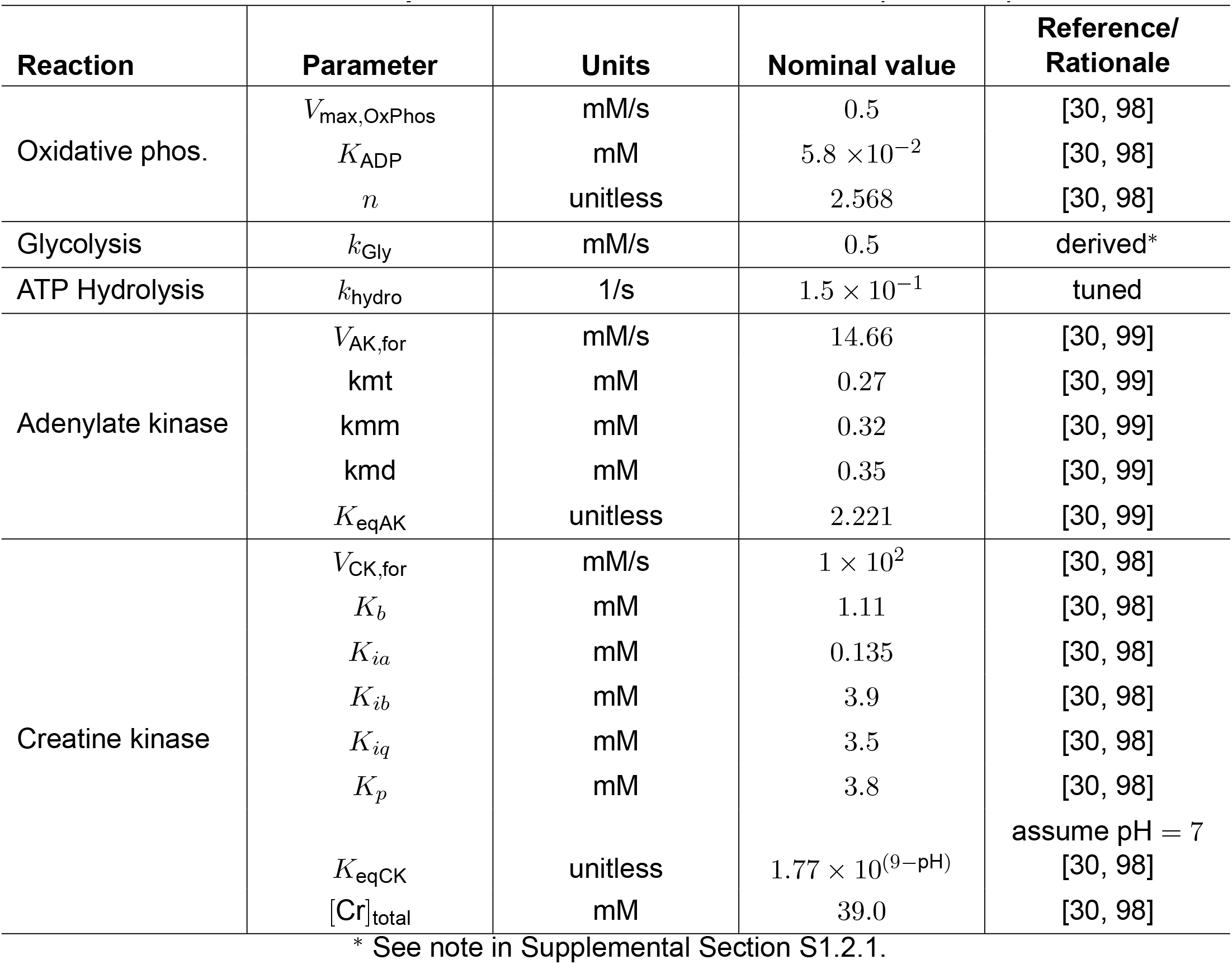
Model parameters for metabolism module (Module 1).

| Reaction | Parameter | Units | Nominal value | Reference/<br>Rationale |
| --- | --- | --- | --- | --- |
| Oxidative phos. | $V_{\max, \text{OxPhos}}$ | mM/s | 0.5 | [30, 98] |
| | $K_{\text{ADP}}$ | mM | $5.8 \times 10^{-2}$ | [30, 98] |
| | $n$ | unitless | 2.568 | [30, 98] |
| Glycolysis | $k_{\text{Gly}}$ | mM/s | 0.5 | derived* |
| ATP Hydrolysis | $k_{\text{hydro}}$ | 1/s | $1.5 \times 10^{-1}$ | tuned |
| Adenylate kinase | $V_{\text{AK,for}}$ | mM/s | 14.66 | [30, 99] |
| | $k_{\text{mt}}$ | mM | 0.27 | [30, 99] |
| | $k_{\text{mm}}$ | mM | 0.32 | [30, 99] |
| | $k_{\text{md}}$ | mM | 0.35 | [30, 99] |
| | $K_{\text{eqAK}}$ | unitless | 2.221 | [30, 99] |
| Creatine kinase | $V_{\text{CK,for}}$ | mM/s | $1 \times 10^2$ | [30, 98] |
| | $K_b$ | mM | 1.11 | [30, 98] |
| | $K_{ia}$ | mM | 0.135 | [30, 98] |
| | $K_{ib}$ | mM | 3.9 | [30, 98] |
| | $K_{iq}$ | mM | 3.5 | [30, 98] |
| | $K_p$ | mM | 3.8 | [30, 98] |
| | $K_{\text{eqCK}}$ | unitless | $1.77 \times 10^{(9-\text{pH})}$ | assume pH = 7<br>[30, 98] |
| | $[\text{Cr}]_{\text{total}}$ | mM | 39.0 | [30, 98] |

\* See note in Supplemental Section S1.2.1.

**Table 6:**
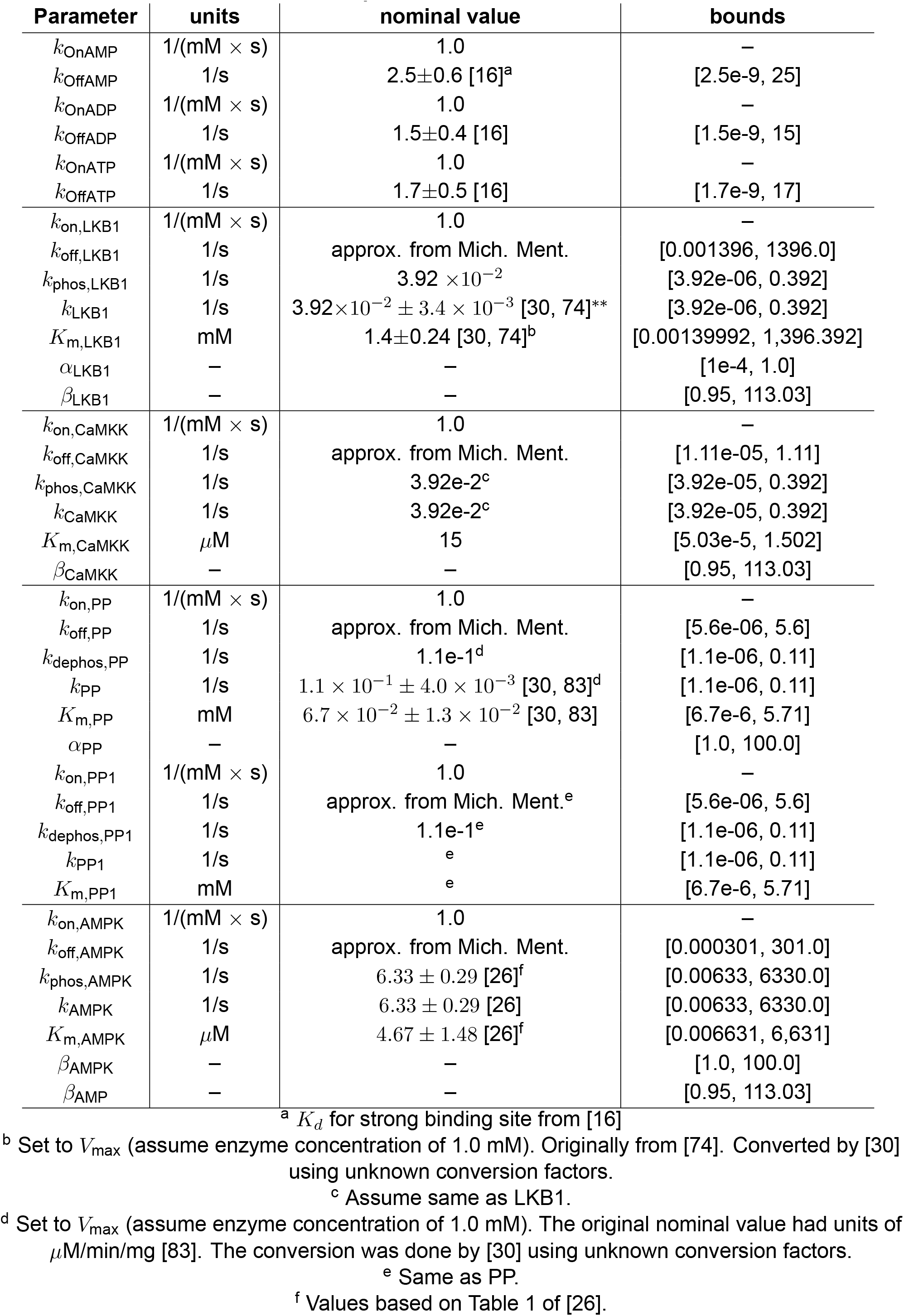
AMPK model parameters. Nominal values and bounds.

| Parameter | units | nominal value | bounds |
| --- | --- | --- | --- |
| $k_{\text{OnAMP}}$ | 1/(mM × s) | 1.0 | – |
| $k_{\text{OffAMP}}$ | 1/s | $2.5 \pm 0.6$ [16]<sup>a</sup> | [2.5e-9, 25] |
| $k_{\text{OnADP}}$ | 1/(mM × s) | 1.0 | – |
| $k_{\text{OffADP}}$ | 1/s | $1.5 \pm 0.4$ [16] | [1.5e-9, 15] |
| $k_{\text{OnATP}}$ | 1/(mM × s) | 1.0 | – |
| $k_{\text{OffATP}}$ | 1/s | $1.7 \pm 0.5$ [16] | [1.7e-9, 17] |
| $k_{\text{on,LKB1}}$ | 1/(mM × s) | 1.0 | – |
| $k_{\text{off,LKB1}}$ | 1/s | approx. from Mich. Ment. | [0.001396, 1396.0] |
| $k_{\text{phos,LKB1}}$ | 1/s | $3.92 \times 10^{-2}$ | [3.92e-06, 0.392] |
| $k_{\text{LKB1}}$ | 1/s | $3.92 \times 10^{-2} \pm 3.4 \times 10^{-3}$ [30, 74]** | [3.92e-06, 0.392] |
| $K_{\text{m,LKB1}}$ | mM | $1.4 \pm 0.24$ [30, 74]<sup>b</sup> | [0.00139992, 1,396.392] |
| $\alpha_{\text{LKB1}}$ | – | – | [1e-4, 1.0] |
| $\beta_{\text{LKB1}}$ | – | – | [0.95, 113.03] |
| $k_{\text{on,CaMKK}}$ | 1/(mM × s) | 1.0 | – |
| $k_{\text{off,CaMKK}}$ | 1/s | approx. from Mich. Ment. | [1.11e-05, 1.11] |
| $k_{\text{phos,CaMKK}}$ | 1/s | $3.92\text{e-}2^{\text{c}}$ | [3.92e-05, 0.392] |
| $k_{\text{CaMKK}}$ | 1/s | $3.92\text{e-}2^{\text{c}}$ | [3.92e-05, 0.392] |
| $K_{\text{m,CaMKK}}$ | $\mu\text{M}$ | 15 | [5.03e-5, 1.502] |
| $\beta_{\text{CaMKK}}$ | – | – | [0.95, 113.03] |
| $k_{\text{on,PP}}$ | 1/(mM × s) | 1.0 | – |
| $k_{\text{off,PP}}$ | 1/s | approx. from Mich. Ment. | [5.6e-06, 5.6] |
| $k_{\text{dephos,PP}}$ | 1/s | $1.1\text{e-}1^{\text{d}}$ | [1.1e-06, 0.11] |
| $k_{\text{PP}}$ | 1/s | $1.1 \times 10^{-1} \pm 4.0 \times 10^{-3}$ [30, 83]<sup>d</sup> | [1.1e-06, 0.11] |
| $K_{\text{m,PP}}$ | mM | $6.7 \times 10^{-2} \pm 1.3 \times 10^{-2}$ [30, 83] | [6.7e-6, 5.71] |
| $\alpha_{\text{PP}}$ | – | – | [1.0, 100.0] |
| $k_{\text{on,PP1}}$ | 1/(mM × s) | 1.0 | – |
| $k_{\text{off,PP1}}$ | 1/s | approx. from Mich. Ment.<sup>e</sup> | [5.6e-06, 5.6] |
| $k_{\text{dephos,PP1}}$ | 1/s | $1.1\text{e-}1^{\text{e}}$ | [1.1e-06, 0.11] |
| $k_{\text{PP1}}$ | 1/s | <sup>e</sup> | [1.1e-06, 0.11] |
| $K_{\text{m,PP1}}$ | mM | <sup>e</sup> | [6.7e-6, 5.71] |
| $k_{\text{on,AMPK}}$ | 1/(mM × s) | 1.0 | – |
| $k_{\text{off,AMPK}}$ | 1/s | approx. from Mich. Ment. | [0.000301, 301.0] |
| $k_{\text{phos,AMPK}}$ | 1/s | $6.33 \pm 0.29$ [26]<sup>f</sup> | [0.00633, 6330.0] |
| $k_{\text{AMPK}}$ | 1/s | $6.33 \pm 0.29$ [26] | [0.00633, 6330.0] |
| $K_{\text{m,AMPK}}$ | $\mu\text{M}$ | $4.67 \pm 1.48$ [26]<sup>f</sup> | [0.006631, 6,631] |
| $\beta_{\text{AMPK}}$ | – | – | [1.0, 100.0] |
| $\beta_{\text{AMP}}$ | – | – | [0.95, 113.03] |

<sup>a</sup> $K_d$ for strong binding site from [16]<sup>b</sup> Set to $V_{\text{max}}$ (assume enzyme concentration of 1.0 mM). Originally from [74]. Converted by [30] using unknown conversion factors.<sup>c</sup> Assume same as LKB1.<sup>d</sup> Set to $V_{\text{max}}$ (assume enzyme concentration of 1.0 mM). The original nominal value had units of $\mu\text{M}/\text{min}/\text{mg}$ [83]. The conversion was done by [30] using unknown conversion factors.<sup>e</sup> Same as PP.<sup>f</sup> Values based on Table 1 of [26].

**Table 7:** Local identifiability.

| Model | Locally Identifiable | NOT Locally Identifiable |
| --- | --- | --- |
| 1 | kOnAMP, kOffAMP, kOnADP, kOffADP, kOnATP, kOffATP, kOnCaMKK, kOffCaMKK, kPhosCaMKK, kOnLKB1, kOffLKB1, kPhosLKB1, kOnPP, kOffPP, kDephosPP, kOnAMPK, kOffAMPK, kPhosAMPK, kOnPP1, kOffPP1, kDephosPP1 |  |
| 2 | kOnAMP, kOffAMP, kOnADP, kOffADP, kOnATP, kOffATP, KmCaMKK, kLKB1, KmLKB1, kPP, KmPP | kCaMKK, kAMPK, KmAMPK, kPP1, KmPP1 |
| 3 | kOnAMP, kOffAMP, kOnADP, kOffADP, kOnATP, kOffATP, kOnCaMKK, kOffCaMKK, kPhosCaMKK, kOnLKB1, kOffLKB1, alphaLKB1, kPhosLKB1, kOnPP, kOffPP, alphaPP, kDephosPP, kOnAMPK, kOffAMPK, kPhosAMPK, betaAMP, kOnPP1, kOffPP1, kDephosPP1 |  |
| 4 | kOnAMP, kOffAMP, kOnADP, kOffADP, kOnATP, kOffATP, KmCaMKK, kLKB1, KmLKB1, alphaLKB1, kPP, KmPP, alphaPP, betaAMP | kCaMKK, kAMPK, KmAMPK, kPP1, KmPP1 |
| 5 | kOnAMP, kOffAMP, kOnADP, kOffADP, kOnATP, kOffATP, kOnCaMKK, kOffCaMKK, kPhosCaMKK, kOnLKB1, kOffLKB1, kPhosLKB1, kOnPP, kOffPP, kDephosPP, kOnAMPK, kOffAMPK, kPhosAMPK, kOnPP1, kOffPP1, kDephosPP1, alphaPP, betaAMP, betaCaMKK, betaLKB1 |  |
| 6 | kOnAMP, kOffAMP, kOnADP, kOffADP, kOnATP, kOffATP, KmCaMKK, kLKB1, KmLKB1, kPP, KmPP, betaAMPK, betaLKB1, betaCaMKK, alphaPP | kCaMKK, kAMPK, KmAMPK, kPP1, KmPP1 |

**Table 8:** Sobol total sensitivity indices for Model 1. Bold values indicate influential parameters with a mean index greater than or equal to 0.01.

| Parameter | Function | max fraction act. AMPKAR |  |  | Time to half-max |  |  |
| --- | --- | --- | --- | --- | --- | --- | --- |
|  |  | WT | LKB1 KO | CaMKK2 KO | WT | LKB1 KO | CaMKK2 KO |
| $k_{\text{OffAMP}}$ | AMP binding | **0.135** ± **0.043** | **0.151** ± **0.044** | **0.151** ± **0.048** | **0.067** ± **0.017** | **0.056** ± **0.012** | **0.073** ± **0.015** |
| $k_{\text{OffADP}}$ | ADP binding | **0.087** ± **0.035** | **0.082** ± **0.034** | **0.085** ± **0.037** | **0.059** ± **0.018** | **0.095** ± **0.033** | **0.056** ± **0.018** |
| $k_{\text{OffATP}}$ | ATP binding | **0.219** ± **0.071** | **0.207** ± **0.059** | **0.184** ± **0.062** | **0.407** ± **0.118** | **0.342** ± **0.095** | **0.359** ± **0.098** |
| $k_{\text{OffCaMKK}}$ | NaN | 0.001 ± 0.001 | **0.010** ± **0.004** | 0.000 ± 0.000 | 0.001 ± 0.001 | **0.019** ± **0.007** | 0.000 ± 0.000 |
| $k_{\text{PhosCaMKK}}$ | AMPK phos. | 0.003 ± 0.003 | **0.084** ± **0.045** | 0.000 ± 0.000 | 0.004 ± 0.002 | **0.155** ± **0.051** | 0.000 ± 0.000 |
| $k_{\text{OffLKB1}}$ | AMPK phos. | 0.001 ± 0.001 | 0.000 ± 0.000 | 0.007 ± 0.003 | 0.005 ± 0.001 | 0.000 ± 0.000 | **0.023** ± **0.010** |
| $k_{\text{PhosLKB1}}$ | AMPK phos. | 0.004 ± 0.003 | 0.000 ± 0.000 | **0.076** ± **0.044** | **0.021** ± **0.008** | 0.000 ± 0.000 | **0.126** ± **0.057** |
| $k_{\text{OffPP}}$ | AMPK dephos. | 0.004 ± 0.003 | 0.005 ± 0.003 | 0.002 ± 0.002 | 0.002 ± 0.001 | 0.004 ± 0.002 | 0.000 ± 0.000 |
| $k_{\text{DephosPP}}$ | AMPK dephos. | 0.003 ± 0.003 | 0.003 ± 0.002 | 0.001 ± 0.001 | 0.001 ± 0.001 | 0.001 ± 0.001 | 0.001 ± 0.000 |
| $k_{\text{OffAMPK}}$ | AMPK kinase act. | **0.183** ± **0.055** | **0.166** ± **0.050** | **0.177** ± **0.053** | **0.108** ± **0.023** | **0.085** ± **0.020** | **0.106** ± **0.028** |
| $k_{\text{PhosAMPK}}$ | AMPK kinase act. | **0.516** ± **0.108** | **0.454** ± **0.084** | **0.480** ± **0.096** | **0.545** ± **0.120** | **0.514** ± **0.122** | **0.493** ± **0.101** |
| $k_{\text{OffPP1}}$ | AMKPAR dephos. | 0.002 ± 0.001 | 0.002 ± 0.001 | 0.002 ± 0.001 | 0.003 ± 0.003 | 0.003 ± 0.003 | 0.002 ± 0.001 |
| $k_{\text{Dephos,PP1}}$ | AMKPAR dephos. | 0.007 ± 0.002 | 0.006 ± 0.001 | 0.007 ± 0.002 | 0.003 ± 0.001 | 0.004 ± 0.002 | 0.003 ± 0.001 |

**Table 9:** Sobol total sensitivity indices for Model 2. Bold values indicate influential parameters with a mean index greater than or equal to 0.01.

| Parameter | Function | max fraction act. AMPKAR |  |  | Time to half-max |  |  |
| --- | --- | --- | --- | --- | --- | --- | --- |
|  |  | WT | LKB1 KO | CaMKK2 KO | WT | LKB1 KO | CaMKK2 KO |
| $k_{\text{OffAMP}}$ | AMP binding | **0.498** ± **0.089** | **0.509** ± **0.098** | **0.724** ± **0.158** | **0.806** ± **0.135** | **0.776** ± **0.131** | **0.478** ± **0.144** |
| $k_{\text{OffADP}}$ | ADP binding | **0.174** ± **0.063** | **0.163** ± **0.054** | **0.038** ± **0.022** | **0.144** ± **0.056** | **0.158** ± **0.062** | **0.218** ± **0.091** |
| $k_{\text{OffATP}}$ | ATP binding | **0.327** ± **0.086** | **0.310** ± **0.081** | **0.155** ± **0.076** | **0.255** ± **0.085** | **0.237** ± **0.088** | **0.242** ± **0.115** |
| $K_{m,\text{CaMKK}}$ | AMPK phos. | **0.127** ± **0.043** | **0.147** ± **0.046** | 0.000 ± 0.000 | **0.149** ± **0.048** | **0.197** ± **0.068** | 0.000 ± 0.000 |
| $k_{\text{LKB1}}$ | AMPK phos. | 0.004 ± 0.003 | 0.000 ± 0.000 | **0.215** ± **0.114** | 0.007 ± 0.004 | 0.000 ± 0.000 | **0.260** ± **0.093** |
| $K_{m,\text{LKB1}}$ | AMPK phos. | **0.011** ± **0.005** | 0.000 ± 0.000 | **0.233** ± **0.079** | **0.018** ± **0.011** | 0.000 ± 0.000 | **0.557** ± **0.286** |
| $k_{\text{PP}}$ | AMPK dephos. | **0.035** ± **0.016** | **0.037** ± **0.015** | **0.012** ± **0.007** | **0.017** ± **0.009** | **0.020** ± **0.011** | **0.085** ± **0.044** |
| $K_{m,\text{PP}}$ | AMPK dephos. | **0.024** ± **0.009** | **0.024** ± **0.010** | 0.009 ± 0.005 | **0.022** ± **0.011** | **0.025** ± **0.014** | **0.036** ± **0.018** |

**Table 10:** Sobol total sensitivity indices for Model 3. Bold values indicate influential parameters with a mean index greater than or equal to 0.01.

| Parameter | Function | max fraction act. AMPKAR |  |  | Time to half-max |  |  |
| --- | --- | --- | --- | --- | --- | --- | --- |
|  |  | WT | LKB1 KO | CaMKK2 KO | WT | LKB1 KO | CaMKK2 KO |
| $k_{\text{OffAMP}}$ | AMP binding | **0.198** ± **0.053** | **0.227** ± **0.064** | **0.208** ± **0.055** | **0.416** ± **0.108** | **0.454** ± **0.133** | **0.415** ± **0.125** |
| $k_{\text{OffADP}}$ | ADP binding | **0.086** ± **0.038** | **0.072** ± **0.040** | **0.081** ± **0.031** | **0.057** ± **0.024** | **0.028** ± **0.012** | **0.063** ± **0.031** |
| $k_{\text{OffATP}}$ | ATP binding | **0.075** ± **0.036** | **0.082** ± **0.046** | **0.074** ± **0.032** | **0.584** ± **0.210** | **0.348** ± **0.145** | **0.517** ± **0.152** |
| $k_{\text{OffCaMKK}}$ | NaN | 0.001 ± 0.000 | 0.007 ± 0.004 | 0.000 ± 0.000 | 0.002 ± 0.001 | **0.023** ± **0.013** | 0.000 ± 0.000 |
| $k_{\text{PhosCaMKK}}$ | AMPK phos. | 0.002 ± 0.001 | **0.055** ± **0.026** | 0.000 ± 0.000 | **0.025** ± **0.027** | **0.248** ± **0.102** | 0.000 ± 0.000 |
| $k_{\text{OffLKB1}}$ | AMPK phos. | 0.002 ± 0.001 | 0.000 ± 0.000 | 0.007 ± 0.005 | 0.010 ± 0.004 | 0.000 ± 0.000 | **0.047** ± **0.052** |
| $k_{\text{PhosLKB1}}$ | AMPK phos. | 0.007 ± 0.004 | 0.000 ± 0.000 | **0.026** ± **0.015** | **0.045** ± **0.023** | 0.000 ± 0.000 | **0.150** ± **0.087** |
| $k_{\text{OffPP}}$ | AMPK dephos. | 0.002 ± 0.002 | 0.004 ± 0.002 | 0.002 ± 0.002 | 0.005 ± 0.005 | **0.016** ± **0.014** | 0.005 ± 0.005 |
| $k_{\text{DephosPP}}$ | AMPK dephos. | 0.001 ± 0.000 | 0.003 ± 0.002 | 0.001 ± 0.000 | 0.002 ± 0.002 | **0.011** ± **0.010** | 0.002 ± 0.002 |
| $k_{\text{OffAMPK}}$ | AMPK kinase act. | **0.036** ± **0.012** | **0.036** ± **0.015** | **0.036** ± **0.013** | **0.052** ± **0.037** | **0.081** ± **0.065** | **0.044** ± **0.029** |
| $k_{\text{PhosAMPK}}$ | AMPK kinase act. | **0.135** ± **0.070** | **0.123** ± **0.075** | **0.134** ± **0.071** | **0.036** ± **0.028** | **0.020** ± **0.014** | **0.034** ± **0.026** |
| $k_{\text{OffPP1}}$ | AMKPAR dephos. | **0.030** ± **0.008** | **0.029** ± **0.010** | **0.031** ± **0.011** | **0.022** ± **0.012** | **0.014** ± **0.009** | **0.023** ± **0.013** |
| $k_{\text{Dephos,PP1}}$ | AMKPAR dephos. | **0.501** ± **0.109** | **0.471** ± **0.128** | **0.508** ± **0.107** | **0.087** ± **0.054** | **0.057** ± **0.048** | **0.072** ± **0.037** |
| $\alpha_{\text{LKB1}}$ | AMPK phos. | 0.001 ± 0.000 | 0.000 ± 0.000 | 0.004 ± 0.004 | 0.004 ± 0.002 | 0.000 ± 0.000 | **0.015** ± **0.009** |
| $\alpha_{\text{PP}}$ | AMPK dephos. | 0.002 ± 0.001 | 0.003 ± 0.002 | 0.002 ± 0.001 | 0.002 ± 0.002 | 0.005 ± 0.003 | 0.002 ± 0.001 |
| $\beta_{\text{AMP}}$ | AMPK kinase act. | **0.106** ± **0.046** | **0.090** ± **0.044** | **0.105** ± **0.050** | **0.056** ± **0.050** | **0.037** ± **0.034** | **0.050** ± **0.038** |

**Table 11:** Sobol total sensitivity indices for Model 4. Bold values indicate influential parameters with a mean index greater than or equal to 0.01.

| Parameter | Function | max fraction act. AMPKAR |  |  | Time to half-max |  |  |
| --- | --- | --- | --- | --- | --- | --- | --- |
|  |  | WT | LKB1 KO | CaMKK2 KO | WT | LKB1 KO | CaMKK2 KO |
| $k_{\text{OffAMP}}$ | AMP binding | **2.38e9 ± 2.79e9** | **2.69e9 ± 2.91e9** | **0.111 ± 0.112** | **0.070 ± 0.055** | **0.044 ± 0.033** | **0.139 ± 0.042** |
| $k_{\text{OffADP}}$ | ADP binding | **0.014 ± 0.010** | **0.014 ± 0.011** | **0.025 ± 0.034** | **0.020 ± 0.022** | 0.006 ± 0.006 | **0.019 ± 0.010** |
| $k_{\text{OffATP}}$ | ATP binding | **0.860 ± 3.147** | **0.852 ± 2.769** | **0.240 ± 0.179** | **0.601 ± 0.980** | **0.604 ± 1.048** | **0.263 ± 0.094** |
| $K_{m,\text{CaMKK}}$ | AMPK phos. | **0.341 ± 0.372** | **0.344 ± 0.343** | 0.000 ± 0.000 | **0.166 ± 0.176** | **0.210 ± 0.165** | 0.000 ± 0.000 |
| $k_{\text{LKB1}}$ | AMPK phos. | 0.000 ± 0.000 | 0.000 ± 0.000 | **0.324 ± 0.220** | **0.039 ± 0.046** | 0.000 ± 0.000 | **0.404 ± 0.113** |
| $K_{m,\text{LKB1}}$ | AMPK phos. | 0.000 ± 0.000 | 0.000 ± 0.000 | **0.223 ± 0.184** | **0.053 ± 0.060** | 0.000 ± 0.000 | **0.184 ± 0.060** |
| $k_{\text{PP}}$ | AMPK dephos. | 0.000 ± 0.000 | 0.000 ± 0.000 | **0.179 ± 0.141** | **0.046 ± 0.053** | 0.000 ± 0.000 | **0.177 ± 0.083** |
| $K_{M,\text{PP}}$ | AMPK dephos. | **0.014 ± 0.010** | **0.014 ± 0.009** | **0.074 ± 0.070** | **0.010 ± 0.005** | 0.007 ± 0.004 | **0.023 ± 0.011** |
| $\alpha_{\text{LKB1}}$ | AMPK phos. | **0.475 ± 0.455** | **0.479 ± 0.412** | **0.107 ± 0.107** | **0.212 ± 0.228** | **0.224 ± 0.254** | **0.051 ± 0.034** |
| $\alpha_{\text{PP}}$ | AMPK dephos. | **0.457 ± 0.431** | **0.461 ± 0.392** | **0.129 ± 0.135** | **0.200 ± 0.206** | **0.204 ± 0.232** | **0.039 ± 0.023** |
| $\beta_{\text{AMP}}$ | AMPK kinase act. | **0.882 ± 0.471** | **0.887 ± 0.416** | **0.385 ± 0.284** | **0.573 ± 0.261** | **0.618 ± 0.289** | **0.206 ± 0.054** |

**Table 12:** Sobol total sensitivity indices for Model 5. Bold values indicate influential parameters with a mean index greater than or equal to 0.01.

| Parameter | Function | max fraction act. AMPKAR |  |  | Time to half-max |  |  |
| --- | --- | --- | --- | --- | --- | --- | --- |
|  |  | WT | LKB1 KO | CaMKK2 KO | WT | LKB1 KO | CaMKK2 KO |
| $k_{OffAMP}$ | AMP binding | **0.092** ± **0.027** | **0.089** ± **0.023** | **0.090** ± **0.026** | **0.181** ± **0.102** | **0.136** ± **0.111** | **0.169** ± **0.081** |
| $k_{OffADP}$ | ADP binding | **0.127** ± **0.042** | **0.054** ± **0.023** | **0.125** ± **0.044** | **0.195** ± **0.146** | **0.082** ± **0.059** | **0.186** ± **0.118** |
| $k_{OffATP}$ | ATP binding | **0.246** ± **0.059** | **0.131** ± **0.048** | **0.249** ± **0.076** | **0.447** ± **0.269** | **0.148** ± **0.100** | **0.425** ± **0.188** |
| $k_{OffCaMKK}$ | NaN | 0.000 ± 0.000 | 0.002 ± 0.001 | 0.000 ± 0.000 | 0.001 ± 0.001 | 0.005 ± 0.003 | 0.000 ± 0.000 |
| $k_{PhosCaMKK}$ | AMPK phos. | 0.002 ± 0.002 | **0.011** ± **0.007** | 0.000 ± 0.000 | 0.004 ± 0.002 | **0.014** ± **0.009** | 0.000 ± 0.000 |
| $k_{OffLKB1}$ | AMPK phos. | 0.003 ± 0.002 | 0.000 ± 0.000 | 0.006 ± 0.003 | **0.014** ± **0.007** | 0.000 ± 0.000 | **0.021** ± **0.010** |
| $k_{PhosLKB1}$ | AMPK phos. | **0.012** ± **0.008** | 0.000 ± 0.000 | **0.023** ± **0.010** | **0.037** ± **0.012** | 0.000 ± 0.000 | **0.060** ± **0.025** |
| $k_{OffPP}$ | AMPK dephos. | **0.036** ± **0.019** | **0.036** ± **0.012** | **0.035** ± **0.018** | **0.457** ± **0.268** | **0.549** ± **0.367** | **0.375** ± **0.219** |
| $k_{DephosPP}$ | AMPK dephos. | **0.026** ± **0.019** | **0.070** ± **0.023** | **0.027** ± **0.020** | **0.084** ± **0.038** | **0.193** ± **0.177** | **0.085** ± **0.049** |
| $k_{OffAMPK}$ | AMPK kinase act. | **0.024** ± **0.013** | **0.031** ± **0.019** | **0.024** ± **0.013** | **0.156** ± **0.081** | **0.183** ± **0.193** | **0.155** ± **0.090** |
| $k_{PhosAMPK}$ | AMPK kinase act. | **0.197** ± **0.058** | **0.118** ± **0.041** | **0.187** ± **0.050** | **0.321** ± **0.115** | **0.061** ± **0.040** | **0.318** ± **0.140** |
| $k_{OffPP1}$ | AMKPAR dephos. | **0.095** ± **0.025** | **0.084** ± **0.024** | **0.090** ± **0.025** | **0.191** ± **0.096** | **0.050** ± **0.052** | **0.196** ± **0.110** |
| $k_{DephosPP1}$ | AMKPAR dephos. | **0.398** ± **0.094** | **0.500** ± **0.098** | **0.387** ± **0.088** | **0.328** ± **0.144** | **0.066** ± **0.049** | **0.311** ± **0.130** |
| $\alpha_{PP}$ | AMPK dephos. | **0.040** ± **0.018** | **0.113** ± **0.041** | **0.043** ± **0.019** | **0.187** ± **0.194** | **0.369** ± **0.232** | **0.143** ± **0.136** |
| $\beta_{AMP}$ | AMPK kinase act. | **0.071** ± **0.030** | **0.046** ± **0.024** | **0.069** ± **0.033** | **0.086** ± **0.041** | **0.012** ± **0.011** | **0.085** ± **0.043** |
| $\beta_{LKB1}$ | AMPK phos. | **0.022** ± **0.011** | 0.010 ± 0.009 | **0.044** ± **0.027** | **0.028** ± **0.023** | 0.004 ± 0.004 | **0.038** ± **0.026** |
| $\beta_{CaMKK}$ | AMPK phos. | 0.000 ± 0.000 | 0.000 ± 0.000 | 0.000 ± 0.000 | 0.000 ± 0.000 | 0.000 ± 0.000 | 0.000 ± 0.000 |

**Table 13:** Sobol total sensitivity indices for Model 6. Bold values indicate influential parameters with a mean index greater than or equal to 0.01.

| Parameter | Function | max fraction act. AMPKAR |  |  | Time to half-max |  |  |
| --- | --- | --- | --- | --- | --- | --- | --- |
|  |  | WT | LKB1<br>KO | CaMKK2<br>KO | WT | LKB1<br>KO | CaMKK2<br>KO |
| $k_{\text{OffAMP}}$ | AMP binding | **0.196** ±<br>**0.071** | **0.261** ±<br>**0.075** | **0.198** ±<br>**0.072** | **0.271** ±<br>**0.060** | **0.401** ±<br>**0.171** | **0.270** ±<br>**0.055** |
| $k_{\text{OffADP}}$ | ADP binding | **0.174** ±<br>**0.080** | **0.075** ±<br>**0.035** | **0.173** ±<br>**0.062** | **0.114** ±<br>**0.037** | **0.076** ±<br>**0.050** | **0.114** ±<br>**0.034** |
| $k_{\text{OffATP}}$ | ATP binding | **0.271** ±<br>**0.084** | **0.139** ±<br>**0.059** | **0.273** ±<br>**0.086** | **0.414** ±<br>**0.126** | **0.243** ±<br>**0.160** | **0.412** ±<br>**0.117** |
| $K_{m,\text{CaMKK}}$ | AMPK phos. | 0.000 ±<br>0.000 | **0.311** ±<br>**0.110** | 0.000 ±<br>0.000 | 0.000 ±<br>0.000 | **0.403** ±<br>**0.249** | 0.000 ±<br>0.000 |
| $k_{\text{LKB1}}$ | AMPK phos. | **0.018** ±<br>**0.029** | 0.000 ±<br>0.000 | **0.020** ±<br>**0.037** | **0.016** ±<br>**0.010** | 0.000 ±<br>0.000 | **0.016** ±<br>**0.010** |
| $K_{m,\text{LKB1}}$ | AMPK phos. | **0.017** ±<br>**0.029** | 0.000 ±<br>0.000 | **0.020** ±<br>**0.037** | 0.006 ±<br>0.007 | 0.000 ±<br>0.000 | 0.006 ±<br>0.007 |
| $k_{\text{PP}}$ | AMPK dephos. | 0.000 ±<br>0.000 | 0.003 ±<br>0.002 | 0.000 ±<br>0.001 | 0.000 ±<br>0.000 | **0.054** ±<br>**0.038** | 0.000 ±<br>0.000 |
| $K_{M,\text{PP}}$ | AMPK dephos. | 0.004 ±<br>0.004 | 0.006 ±<br>0.003 | 0.003 ±<br>0.004 | 0.001 ±<br>0.001 | **0.233** ±<br>**0.116** | 0.001 ±<br>0.001 |
| $\alpha_{\text{PP}}$ | AMPK dephos. | **0.598** ±<br>**0.127** | **0.271** ±<br>**0.108** | **0.595** ±<br>**0.130** | **0.595** ±<br>**0.110** | **0.250** ±<br>**0.161** | **0.595** ±<br>**0.097** |
| $\beta_{\text{AMP}}$ | AMPK kinase act. | 0.001 ±<br>0.001 | 0.000 ±<br>0.000 | 0.001 ±<br>0.001 | 0.002 ±<br>0.002 | 0.000 ±<br>0.000 | 0.002 ±<br>0.002 |
| $\beta_{\text{LKB1}}$ | AMPK phos. | 0.000 ±<br>0.000 | **0.258** ±<br>**0.095** | 0.000 ±<br>0.000 | 0.000 ±<br>0.000 | **0.392** ±<br>**0.237** | 0.000 ±<br>0.000 |
| $\beta_{\text{CaMKK}}$ | AMPK phos. | 0.000 ±<br>0.000 | 0.000 ±<br>0.000 | 0.000 ±<br>0.000 | 0.000 ±<br>0.000 | **0.012** ±<br>**0.007** | 0.000 ±<br>0.000 |

